# Engineering transplantable jejunal mucosal grafts using primary patient-derived organoids from children with intestinal failure

**DOI:** 10.1101/854083

**Authors:** Laween Meran, Isobel Massie, Anne Weston, Riana Gaifulina, Peter Faull, Michael Orford, Anna Kucharska, Anna Baulies, Elizabeth Hirst, Julia Konig, Alfonso Maria Tedeschi, Alessandro Filippo Pellegata, Susanna Eli, Ambrosius P. Snijders, Lucy Collinson, Nikhil Thapar, Geriant Thomas, Simon Eaton, Paola Bonfanti, Paolo De Coppi, Vivian S.W. Li

## Abstract

Intestinal failure (IF), following extensive anatomical or functional loss of small intestine (SI), has debilitating long-term effects on infants and children with this condition. Priority of care is to increase the child’s length of functional intestine, jejunum in particular, to improve nutritional independence. Here we report a robust protocol for reconstruction of autologous intestinal mucosal grafts using primary IF patient materials. Human jejunal intestinal organoids derived from paediatric IF patients can be expanded efficiently *in vitro* with region-specific markers preserved after long-term culture. Decellularized human intestinal matrix with intact ultrastructure is used as biological scaffolds. Proteomic and Raman spectroscopic analyses reveal highly analogous biochemical composition of decellularized human SI and colon matrix, implying that they can both be utilised as scaffolds for jejunal graft reconstruction. Indeed, seeding of primary human jejunal organoids to either SI or colonic scaffolds *in vitro* can efficiently reconstruct functional jejunal grafts with persistent disaccharidase activity as early as 4 days after seeding, which can further survive and mature after transplantation *in vivo*. Our findings pave the way towards regenerative medicine for IF patients.

## INTRODUCTION

Infants with intestinal failure (IF) have a reduction in functional intestinal mass below the minimal requirement to satisfy nutrient and fluid needs to sustain growth^1^. This may be caused by anatomical loss due to short bowel syndrome (SBS), dysmotility due to neuromuscular intestinal diseases, or congenital epithelial defects of the intestine^2^. IF patients may become dependent on intravenous feeding known as parenteral nutrition (PN), which is associated with numerous complications including bacterial overgrowth, line sepsis, central venous access thrombosis or PN-related liver disease^3,4^. Ultimately, children with irreversible IF are referred for small bowel transplantation. However, many children die on the transplant waiting list due to shortage of suitable organs. Among those who receive a transplant, patient survival at 5 years is only 58% due to sepsis, graft failure and complications of long-term immunosuppression^5^. Given the high morbidity and mortality associated with current treatment for IF, there is an urgent unmet clinical need for research into novel treatment strategies for these patients.

Autologous tissue engineering of functional intestinal grafts, through combination of biomaterials and patient-derived cells, presents an innovative alternative approach to treat IF patients^6^. Multiple studies have previously reported the pre-clinical developments of tissue engineering methodologies in other organs such as oesophageal, skeletal muscle, liver and lung reconstruction^7-13^. However, while engineering of simpler tissues such as skin and cornea are established in clinical practice^14,15^ examples of successful clinical applications of more complex organs have only been demonstrated in few case reports of airway and bladder reconstruction^16,17^. To date there have been no clinical studies of bioengineered small intestinal grafts in humans. Intestinal reconstruction using patient-derived cells would negate the need for immunosuppression, which would circumvent the complications of graft-host rejection, and risk of infection and cancer^5^. Furthermore, tissue engineering strategies to IF should enable personalised grafts whereby length, and cellular or scaffold composition may be modified based on the individual patient’s condition. For instance, SBS is a consequence of massive full thickness anatomical loss of the small intestine, leading to inadequate enteral absorption^3,18-20^. The most predominant pathologies of paediatric SBS include necrotizing enterocolitis, intestinal atresia, gastroschisis, malrotation with volvulus and Crohn’s disease^4^, which require reconstruction of a full thickness intestinal wall graft (including mucosa, submucosa and muscularis layers) to fit infant’s dimensions. On the other hand, patients with neuromuscular intestinal diseases such as extensive Hirschsprung’s disease usually have a healthy intestinal epithelium but dysfunctional neuromuscular wall, where reconstruction of a neuromuscular graft capable of peristalsis is paramount^21^. Conversely, patients with congenital epithelial defects such as Microvillus Inclusion Disease retain intact neuromuscular function^22^. Thus, reconstruction of a purely mucosal graft with gene correction would be most beneficial to these patients.

The success of tissue engineering requires optimal source of cells and scaffolds^23^. Intestinal cells could be derived from human induced pluripotent stem cells (hiPSCs), human embryonic stem cells (hESCs) or primary human somatic intestinal stem cells (hISCs). To achieve autologous transplantation, patient-derived hiPSCs and hISCs are the preferred options. While iPSCs have high potential to regenerate all the required cell sources needed for full thickness intestinal reconstruction, clinical use on paediatric patients remains controversial due to the potential risk of chromosomal aberrations and teratoma development during the reprogramming process^24-28^. Recent advances on the establishment of 3-dimensional human intestinal organoid (hIO) culture system has revolutionised the field of regenerative medicine^29,30^. These mini-hIO culture enables unlimited expansion of primary hISCs while maintaining multipotency and genetic stability. They are seemingly safer and ethically acceptable alternative cell source than hiPSCs for clinical use. However, the efficiency of establishing primary human small intestinal organoids derived from IF patients with limited starting materials has never been demonstrated.

hISCs could be delivered using biodegradable materials such as polyglycolic acid/poly-L-lactic acid polymers^31-33^, however the lack of fine microarchitectural details as well as biological cues responsible for cell engraftment and self-organisation have so far largely limited their translation. On the other hand, biological extracellular matrix (ECM) obtained from decellularization of native organs represent a more physiological alternative to synthetic scaffolds^7,10,34,35^.

Here, we report a robust protocol to generate and expand hIOs derived from biopsies of paediatric IF patients undergoing endoscopy. We further optimise the decellularization protocol to native human intestinal tissues obtained from paediatric patients undergoing intestinal resections and perform in-depth characterisation of the scaffold structure and composition. We focus on reconstruction of a functional jejunal graft, where majority of food digestion and absorption occurs and is crucial to restore nutritional autonomy in IF patients. Our results show that the engineered jejunal grafts can differentiate *in vitro* with digestion and absorption functions, which can further survive and form intestinal lumens *in vivo*. The current findings provide proof-of-concept to engineer autologous, functional and transplantable jejunal mucosal grafts derived from IF patients, and pave the way towards the ultimate reconstruction of full thickness intestinal graft for children with IF and SBS.

## RESULTS

### Robust generation and expansion of human small intestinal organoids derived from paediatric IF patients

The primary challenge of engineering autologous grafts for IF patients is to efficiently isolate and expand patient-derived intestinal epithelial cells from very limited starting materials such as endoscopic biopsies. To optimise the protocol, a maximum of two endoscopic intestinal epithelial biopsies (∼2mm size) from paediatric patients were used for crypt isolation and organoid generation (Fig. 1a). In total, organoids were established from 11 individuals, among which 5 were diagnosed with IF clinically (Supplementary Table 1). On average, we were able to establish 3 to 5 organoid units from two endoscopic biopsies by week 4, which can be further expanded to over a million cells by week 8 for scaffold seeding (Fig. 1a-b). The expansion efficiency was similar in all three regions (duodenum, jejunum or ileum) of the small intestine (Fig. 1c). Quantitative reverse transcription-PCR (qRT-PCR) showed that the organoids expressed region-specific functional markers: the apical brush border enzyme cytochrome b reductase 1 (*CYBRD1*) and the iron transporter solute carrier family 40 member 1 (*SLC40A1*) in duodenal organoids; the brush border digestive enzymes sucrase isomaltase (*SI*) and lactase (*LCT*) in jejunal organoids; and the apical bile acid transporter (*SLC10A2*) and the basolateral organic solute transporter (*OSTB*) in ileal organoids (Fig. 1d). We focused on the expansion and maintenance of jejunal organoids for subsequent graft reconstruction experiments as 90% of digestion and absorption of significant macro- and micro-nutrients occur in the proximal 100-150cm of the jejunum^36^. Long-term culture of the jejunal organoids after significant passaging time (P > 25) were able to retain the expression of the region-specific functional markers, indicating that the organoids have cell intrinsic programme to retain their regional identities after prolonged culture (Fig. 1e).

**Figure 1.**
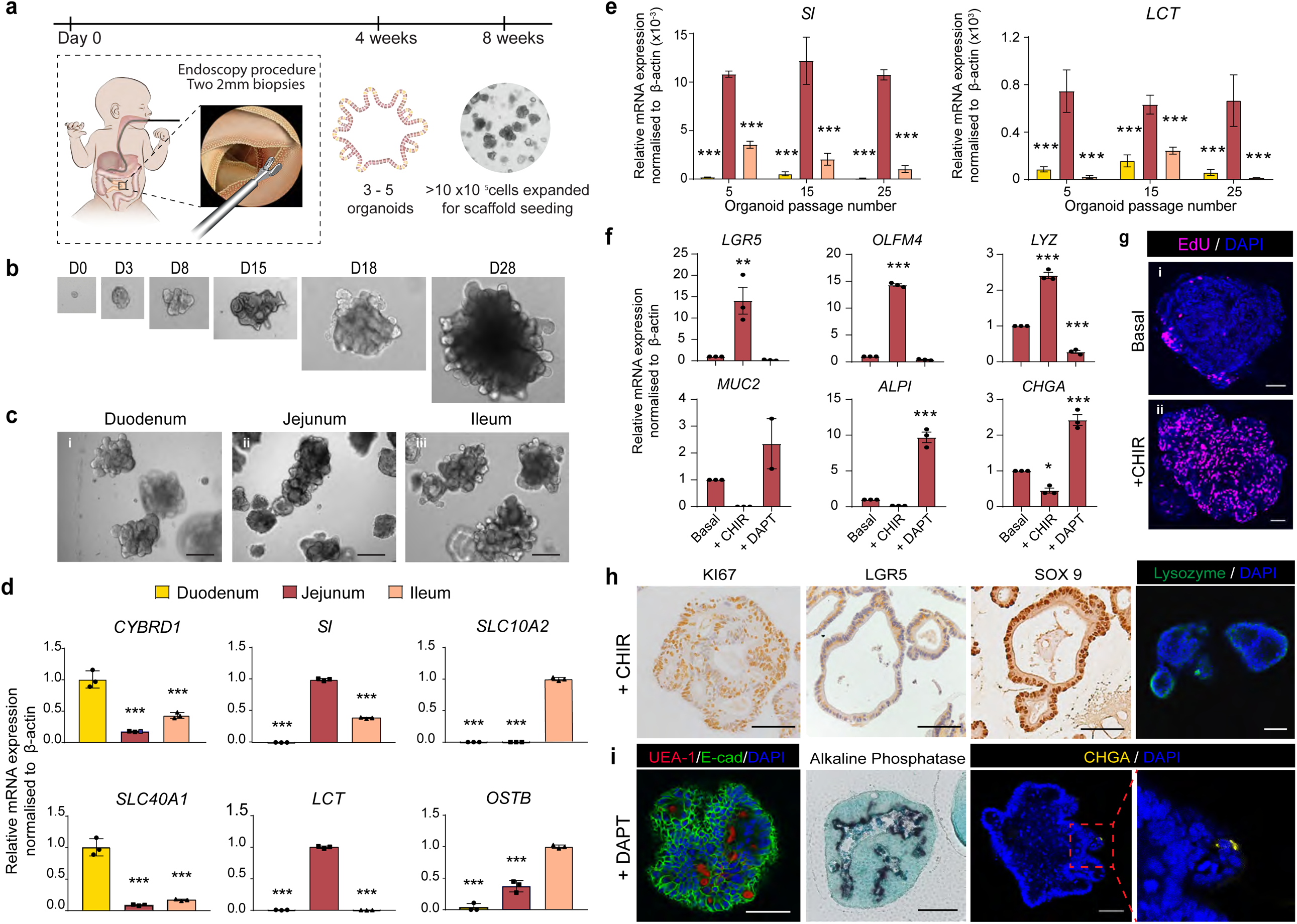
Generation and characterisation of primary intestinal organoids derived from targeted paediatric patient group. (a) Schematic overview demonstrating the expansion timeline when harvesting intestinal crypts endoscopically from paediatric patients. (b) Time course cultures of paediatric small intestinal organoids. (c) First passage expansion of duodenal (i), jejunal (ii) and ileal organoids (iii). Scale bars represent 200μm. (d) Quantitative RT-qPCR analysis of human duodenal, jejunal and ileal organoids for functional duodenal markers (CYBRD1; SLC40A1), jejunal markers (SI; LCT) and ileal markers (SLC10A2; OSTB). Data represent mean ± s.e.m., the experiment was repeated three times. ***P<0.001, two-way ANOVA. (e) Quantitative RT-qPCR analysis of human jejunal organoids at passages 5, 15 and 25 for jejunal specific markers SI and LCT. Data represent mean ± s.e.m. (n=3). ***P < 0.001, two-way ANOVA. (f) Quantitative RT-qPCR analysis of human jejunal organoids treated with GSK3β inhibitor chir99021 (+CHIR), Notch inhibitor (+DAPT). Data represent mean ± s.e.m. (n=3). The experiment was performed three times. *P < 0.5, **P < 0.01, ***P < 0.001, two-way ANOVA. (g) Representative images of EdU incorporation staining of human jejunal organoids in basal culture condition (i) and expansion condition (+CHIR) (ii). Scale bars represent 30μm. (h,i) Representative immunostaining images of human jejunal organoids cultured in expansion conditions (h) or differentiation condition (i) using the indicated antibodies. Scale bars represent 100μm.

Next, we tested the proliferation and differentiation potential of these jejunal organoids after prolonged expansion culture. In addition to the previously published human intestinal organoid media^29^, we further treated the organoids with GSK3β inhibitor (CHIR99021) to boost Wnt signalling and promote proliferation, or with gamma-secretase inhibitor (DAPT) to inhibit Notch signalling and drive differentiation. qRT-PCR analysis showed that stem cell (*OLFM4* and *LGR5*) and Paneth cell (*LYZ*) genes were significantly upregulated while differentiation genes (*MUC2, ALPI* and *CHGA*) were downregulated upon CHIR-treatment (Fig. 1f). A strong induction of proliferation was also observed in CHIR-treated organoids, which was accompanied by protein expression of proliferation (Ki67), stem cell (LGR5, SOX9) and Paneth cell (LYZ) markers (Fig. 1g-h). Conversely, DAPT-treated organoids displayed loss of stem cell and Paneth cell markers and gain of differentiation markers in both mRNA and protein levels (Fig. 1f, i), indicating that the differentiation capacity of these organoids was not affected by the prolonged expansion. Together, the results demonstrate that IF patient-derived organoids can be generated robustly from as little as two 2mm-biopsies, they multiply rapidly *ex vivo* under expansion media (+CHIR) while maintaining their initial regional identity and differentiation potential upon Notch inhibition (+DAPT).

### Decellularization of human small intestinal and colonic ECM scaffolds

Previous studies have described decellularization protocols to generate rodent or piglet small intestinal (SI) ECM scaffolds^34,37^. A more recent study has further attempted to fabricate human SI ECM scaffolds but failed to demonstrate successful decellularization^38^. Here we optimise the decellularization protocols for both human SI and colonic ECM scaffolds and compare their structural and biochemical compositions. Native human SI and colonic tissue was collected from paediatric patients undergoing intestinal resection (Supplementary Table 1), where circumferential intestinal adipose tissue was first dissected and removed from the serosal layer. Excess clinical intestinal tissues are often collected without intact mesentery, thus cannot be decellularized via perfusion. Instead, the native tissues were decellularized using detergent-enzymatic treatment (DET) via a series of immersion and agitation to remove all cellular components while preserving the microarchitecture. Indeed, histology analysis demonstrated absence of cellular and nuclei contents as revealed by H&E and DAPI staining in the decellularized scaffolds, while the intestinal crypt-villus axis were well preserved (Fig. 2a,b). Immunofluorescent staining confirmed the presence of the key ECM protein collagen in both SI and colonic decellularized scaffolds (Fig. 2a). The ultrastructure of the ECM scaffolds was further examined using scanning electron microscopy, which showed remarkable preservation of the microarchitecture of mucosa, submucosa and muscularis layers. Importantly, intact crypt-villus axis of the SI scaffold and crypts of the colonic scaffold in the mucosal layer was clearly identified after decellularization (Fig. 2c), which offer ideal natural biological platforms for organ reconstruction. In addition, we have also optimised the decellularization protocol of piglet SI confirming the removal of the nuclei and the remnants of the ECM component (Supplementary Fig. 1a,b) In particular, the decellularised tissue did not differ substantially from the fresh intestine in protein content, Collagen I, Collagen IV, Fibronectin, Laminin and macro-structure (Supplementary Fig. 1c,d).

**Figure 2.**
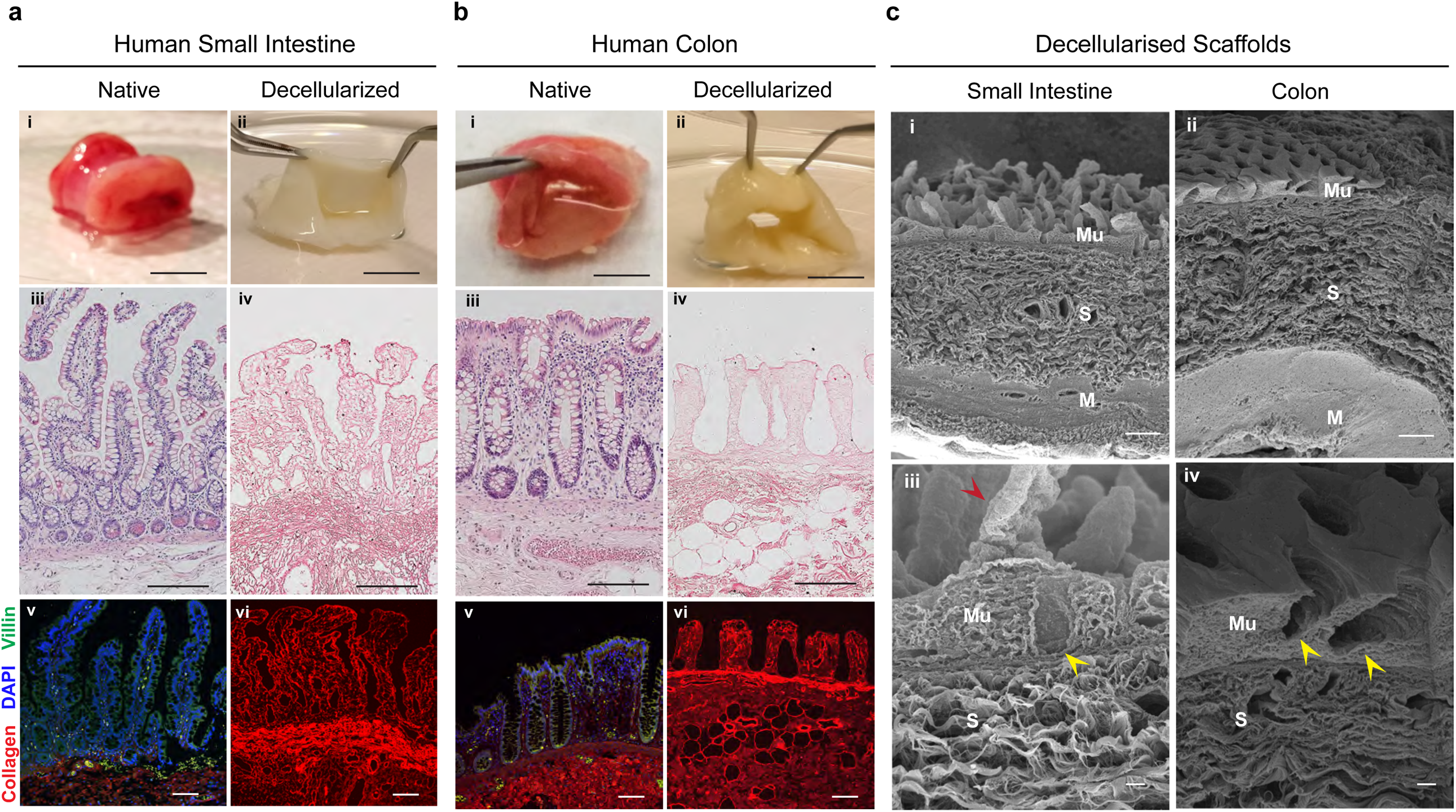
Decellularization of human small intestinal and colon scaffolds. (a,b) Representative images of native human SI (a) and colon (b) before (i) and after decellularization respectively (ii). Scale bars represent 1cm. Representative H&E histological images showing intestinal tissues before (iii) and after decellularization (iv) in both SI (a) and colon (b). Scale bars represent 200μm. Representative images of immunofluorescent staining of intestinal tissues before (iii) and after decellularization (iv) in both SI (a) and colon (b) using the indicated antibodies. Scale bars represent 100μm. (c) Representative images of scanning electron microscopy analysis of the SI (i and iii) and colonic scaffold (ii and iv) highlighting microarchitecture of the mucosa (Mu) submucosa (S) and muscularis (M). Yellow arrow heads indicate intestinal crypts. Red arrow heads indicate villi structure present on the SI scaffold. Scale bars represent 100μm (i and ii) and 10μm (iii and iv).

### Compositional analyses of human small intestine and colon scaffolds

Next, we examined if the biomolecular composition of the scaffolds is also preserved after decellularization. Raman spectroscopy was used to compare the spectra profiles of the native tissues and their corresponding decellularized scaffolds. The overall spectra profiles between native and decellularized scaffolds were highly analogous, indicating that the gross biomolecular composition was preserved (Fig. 3a). On the other hand, a handful of differential spectra was noted across the three tissue layers (mucosa, submucosa and muscle). For instance, the distinct nucleic acid peaks at 726cm^-1^, 780cm^-1^ and 1575cm-1 wavenumber - indicative of the ring breathing mode of nucleotides - were detected preferentially in the native SI and colonic mucosa with dense cellular populations, where the peaks were either significantly reduced or absent in the decellularized scaffolds. Consistently, characteristic lipid peaks at 1078cm-1 and 1303cm^-1^ - indicative of the ν(C-C) skeletal of acyl backbone of lipids and methylene bending mode - was also noted to be more intense in both native mucosa and muscle spectra, which could be associated with the lipid-rich cellular membranes of these two cell-enriched layers. Conversely, the spectral peak at 570cm^-1^ was strongly enriched in both decellularized SI and colonic scaffolds. This peak has previously been attributed to CO_2_ rocking and the S-S bridge of cysteine, proline and tryptophan that are likely the characteristics of the ECM^39^. Similarly, the peak at 1418cm^-1^, which is attributed to CH_2_ bending mode of proteins and lipids, was also detected in the decellularized scaffolds. The enrichment of these two peaks in the decellularized scaffolds highlights the unique properties of the structural ECM scaffolds that lack a dense cell population. Interestingly, minimal difference in peak intensity of the spectra profile was observed between native and decellularized scaffolds in the submucosal region that contains the least cellular density, suggesting that the main difference between the native and decellularized tissues is likely due to the removal of the cellular mass rather than the DET process itself. Of note, the overall spectra profiles between SI and colonic scaffolds were highly analogous, suggesting that the biochemical composition is largely conserved between SI and colonic scaffolds.

**Figure 3.**
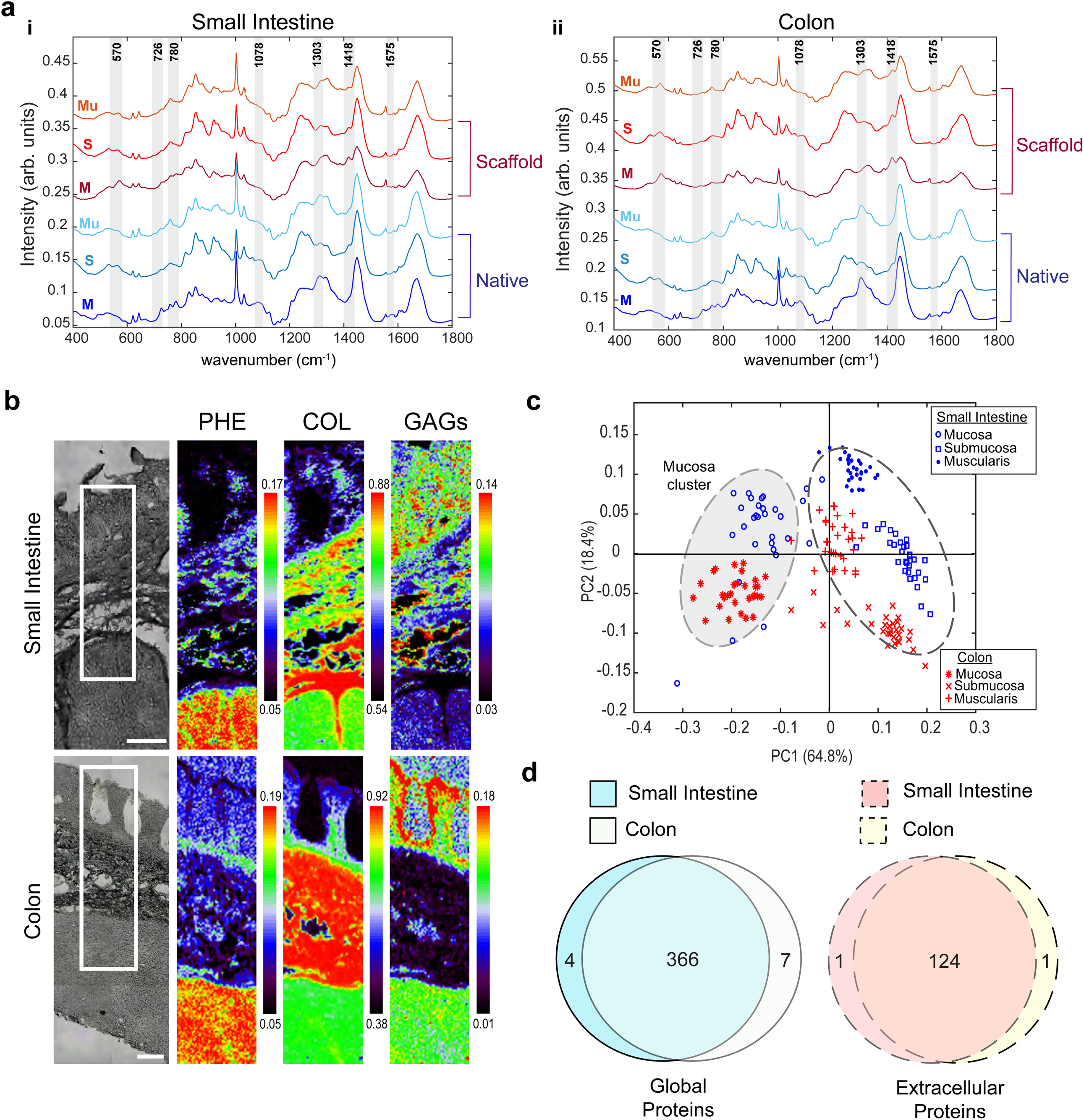
Characterisation of human decellularized intestinal scaffolds by Raman spectroscopy and mass spectrometry. (a) Average Raman spectra from comparable histological regions of the native tissue (blue lines) and decellularized scaffolds (red lines) of SI (i) and colon (ii) samples. (b) Multivariate curve resolution - alternating lease squares false coloured heat map analysis of Phenylalanine (PHE), Collagen (COL) and Glycosaminoglycans (GAGs) in the human SI and colon scaffolds using Raman microspectroscopy. (c) Principal component analysis score plot used to differentiate the Raman spectra generated from each histologically distinct segments of the small intestine scaffolds (blue) and colonic scaffolds (red). (d) Venn diagrams showing the comparative global total proteins and extracellular proteins detected in the SI and colonic scaffolds by mass spectrometry. Data represents samples from 4 biologically independent patient samples in each group.

To demonstrate the spatial distribution of intestinal ECM components in the decellularized scaffolds, Raman spectroscopy was further used for grouping spectra with a similar profile and visualising their spatial localisation via Raman heatmaps. In particular, we noted that collagen was mainly localised to the submucosal layer, while phenylalanine - indicating most proteinaceous regions - was most abundant in the muscularis propria (Fig. 3b). On the other hand, glycosaminoglycans (GAGs), indicated by glucosamines spectra, were highly enriched in the mucosal layer (Fig. 3b). Consistent to the spectra profiling, the ECM component distributions were also highly similar between SI and colonic scaffolds. The results highlight the distinct biomolecular compositions for each histological layer of the scaffolds after decellularization process, which offer essential structural and biochemical cues for cellular reconstitution. Principal component analysis of the spectra profiles readily segregated the spectra into distinct clusters based on their layer identities (Fig. 3c). Remarkably, the mucosal spectra of both SI and colonic scaffold was tightly clustered together, which was distinct from the submucosal and muscularis spectra which are positively characterised (Fig. 3c and Supplementary Fig. 2a). The data suggest that the biochemical composition of the mucosal layer from both SI and colon are more similar to each other than their corresponding deeper histological layers of the scaffolds.

To further characterise the biomolecular profiles, mass spectrometry was used to generate a global proteomic profile of the decellularized SI and colonic ECM scaffolds (n=4 each). This revealed a total of 377 proteins detected in the scaffolds, among which 126 were ECM proteins (Supplementary Table 2). Strikingly, majority of the proteins were detected in both SI and colonic scaffolds, while only 11/377 total proteins and 2/126 ECM proteins were detectable in either SI or colonic scaffolds (Fig. 3d). These included 17 collagen subtypes and 5 Laminin subtypes (Supplementary Table 3). Among the 2 ECM proteins, Defensin-5A (DEFA5) was detectable only in SI scaffold while Thrombospondin-4 (THBS4) was detectably only in colonic scaffolds. Immunohistochemical staining confirmed the restricted expression of DEFA5 in SI but not colon, which was secreted by SI-specific Paneth cells (Supplementary Fig. 2b). THBS4, on the other hand, was expressed in both native SI and colon (Supplementary Fig. 2c), suggesting that the difference noted in mass spectrometry might be due to the detection limit in the SI ECM scaffolds rather than actual expression difference. Altogether, comprehensive analysis of the proteomic profiles of the decellularized intestinal scaffolds demonstrate the highly analogous biochemical composition between human SI and colonic matrix, implicating that both can potentially be used for jejunal graft reconstruction.

### *In vitro* reconstruction of jejunal mucosal grafts

To enhance the bioengineering efficiency, we further generated primary human intestinal fibroblasts isolated from the same intestinal tissues used to derive organoids or SI scaffolds (Supplementary Table 1). Primary human intestinal fibroblasts could be cultured and expanded for up to 10 passages while retaining strong expression of typical stromal matrix markers fibronectin, vimentin, fibroblast surface protein marker-1, laminin α5 and with scattered weaker co-expression of αSMA (Supplementary Fig. 3). This indicates a mixed population of intestinal fibroblasts and myofibroblasts, which are both essential stromal niche for the survival and maintenance of hISCs. To reconstruct a jejunal graft, primary human intestinal fibroblasts were first injected into the scaffolds at the mucosal-submucosal boundary and maintained in static culture for 3 days to recreate the native microenvironment (Fig. 4a). Primary human jejunal organoids were subsequently seeded onto the luminal surface of the scaffold, which was mounted on a customised scaffold holder. The mounted graft was maintained under static conditions for another 4 days, which was then transferred to a dynamic culture condition using a perfusion bioreactor (Fig. 4a). The grafts were maintained under dynamic conditions for at least 14 days before *in vivo* transplantation or histological analysis.

**Figure 4.**
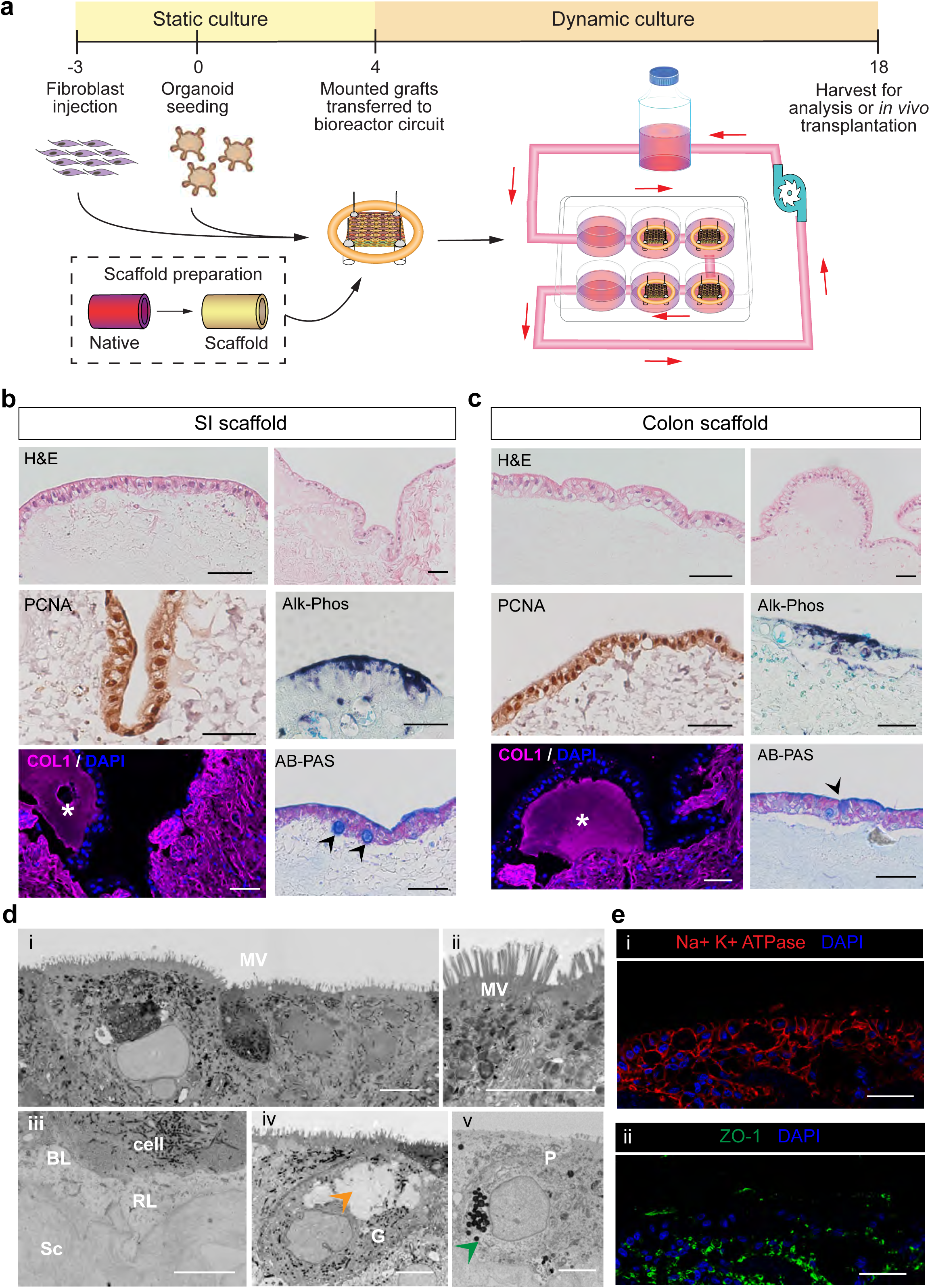
Bioengineering of the human jejunal mucosal graft *in vitro*. (a) Schematic outline of the scaffold seeding strategies under static and dynamic culture using the indicated bioreactor circuit. (b,c) Representative histological images of the jejunal grafts reconstructed using human SI scaffolds (b) or human colonic scaffolds (c) after 14 days of dynamic culture using the indicating staining methods or antibodies. New matrix deposition is shown by newly synthesised collagen (white asterisks). (d) Representative images of electron microscopy analysis of the jejunal constructs showing microvilli (MV) (i and ii); basement membrane with basal lamina (BL) and reticular lamina (RL) at the scaffold (Sc) border (iii); Goblet cell (G) with mucous vesicles indicated by the orange arrow head (iv) and Paneth cell (P) with secretory vesicles indicated by the green arrow head (v). (e) Representative immunofluorescent images of the jejunal grafts seeded on human small intestine scaffolds using the indicated antibodies. All scale bars represent 50μm.

Human jejunal organoids were seeded on piglet SI scaffolds for initial optimisation. Micro-CT imaging was performed on the seeded scaffolds to assess the volume and distribution of epithelial cells on the scaffold, which showed a full surface area coverage of the scaffold (Supplementary Fig. 4a). Histology analysis of the reconstructed grafts showed an organised monolayer of columnar cells on the scaffold (Supplementary Fig. 4b). Regions of collagen-positive new matrix deposition were detected with structures resembling new villi (Supplementary Fig. 4b,c). In addition, all intestinal cell types were readily detected in the graft, including lysozyme-positive Paneth cells, AB-PAS-positive goblet cells and alkaline phosphatase/ALPI-positive enterocytes (Supplementary Fig. 4d-f). Importantly, the jejunal-specific enzyme sucrase isomaltase was also broadly detected on the brush boarder of the epithelial cells, indicating that the region-specific identity was preserved in the engineered graft (Supplementary Fig. 4g). Proliferation marker Ki67 was also detected in the graft (Supplementary Fig. 4h).

Next, we examined the graft reconstruction and differentiation potential by seeding human jejunal organoids on human SI or colonic scaffolds. Similar to piglet scaffold data, histology analysis revealed a continuous monolayer of columnar epithelial cells distributed evenly along the decellularized human SI scaffold surfaces (Fig. 4b). Immunostaining confirmed that proliferation (as indicated by PCNA) and differentiation (ALPI to mark enterocytes and AB-PAS to mark goblet cells) potential was maintained in the reconstructed grafts. Electron microscopy analysis further demonstrated the presence of microvilli (brush boarder of enterocytes), basement membrane (as indicated by basal lamina and reticular lamina features between the scaffold and the epithelial cells), goblet cell mucous vesicles and Paneth cell secretory vesicles (Fig. 4d). Tight junction (ZO1) and polarity (Na+/K+/ATPase) markers were also expressed on the reconstructed jejunal scaffold (Fig. 4e). The presence of basement membrane and junction markers is encouraging as they are essential for the barrier function of the intestine. Interestingly, immunofluorescent staining of collagen showed new matrix deposition in multiple regions underneath the epithelial cells, suggesting ECM remodelling mediated by epithelial cells (Fig. 4b). Similar jejunal graft reconstruction results were observed using human colonic ECM scaffold, where proliferation, differentiation and ECM remodelling were all detected (Fig. 4c). Importantly, the SI-specific expression of ALPI was also detected on the seeded colonic scaffold (Fig. 4c), suggesting that both human SI and colonic ECM scaffolds can be used as biological platforms for reconstruction of jejunal grafts with full proliferation and differentiation potential. Together, the results indicate that human jejunal grafts can be robustly regenerated using either piglet, human SI or colonic scaffolds with capacity for absorptive, digestive and barrier function.

### *Ex vivo* functional analysis of the engineered jejunal grafts

Next, we examined the absorptive and digestive function of the bioengineered grafts at different time points of culture (Fig. 5a). First, the absorptive capacity of the reconstructed jejunal grafts was tested at day 18 of culture by adding the fluorescently labelled peptide β-Ala-Lys-AMCA to the culture. Immunofluorescent staining confirmed the presence of β-AMCA peptides in the epithelial cells, indicating that the engineered graft possesses peptide absorptive function (Fig. 5b). Plasma citrulline levels positively correlate to enterocyte mass and absorptive function and are used as a clinical biomarker of intestinal failure^40^. Citrulline levels were measured in the bioreactor circuit at day 0, day 11 and day 25 of culture as a surrogate marker of enterocyte growth on the scaffolds. There was a continuous increase in citrulline concentrations from day 0 to day 25 in the jejunal grafts regenerated in all three scaffold types, implicating the formation of healthy functional enterocytes in all constructs (Fig. 5c). To further demonstrate digestive function of the reconstructed jejunal grafts, we examined the jejunal enterocyte brush border enzyme sucrase isomaltase function by challenging the constructs with sucrose and measuring the glucose (one of the metabolic products) production at various time point of the culture. Remarkably, glucose production was detected as early as day 4 culture in all three scaffold types, indicating that the disaccharidase function of the jejunal grafts are present throughout the *ex vivo* culture (Fig. 5d). The findings confirmed the presence of absorptive and digestive functions of the engineered human jejunal grafts regenerated on all ECM scaffold types.

**Figure 5.**
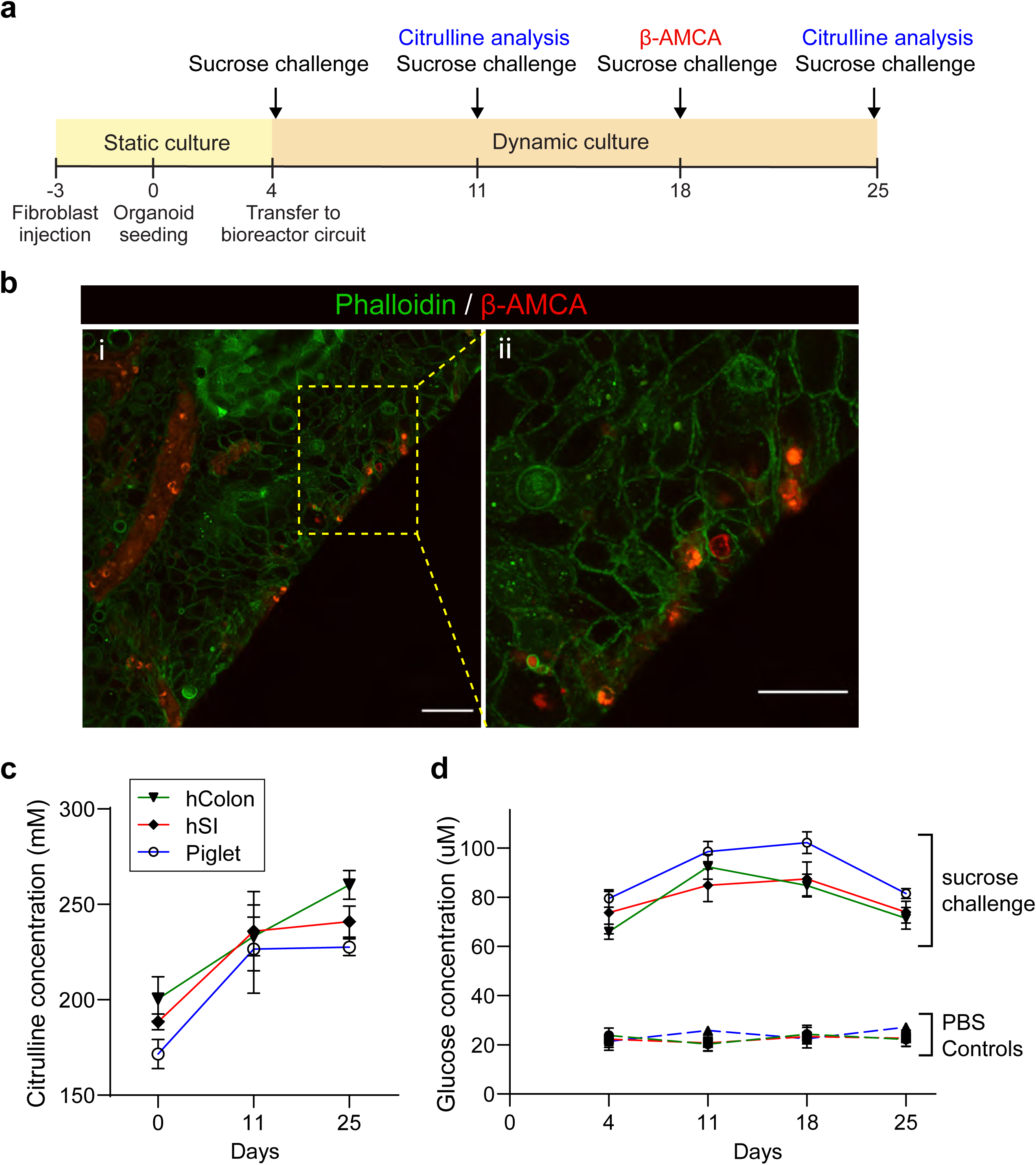
Functional assessment of the absorptive and digestive capacity of the engineered jejunal graft. (a) Experimental design for the functional analysis of the engineered jejunal grafts. (b) Immunofluorescent staining showing β-AMCA peptide (red) uptake on the jejunal grafts. Phalloidin staining (green) indicates epithelial cell boundaries. High magnification image is shown in (ii). Scale bars represent 30μm. (c) Supernatant of the corresponding jejunal graft culture was collected at days 0, 11 and 25 after organoids seeding for measurement of citrulline concentration. (d) Glucose concentration was measured following a sucrose challenge delivered to each scaffold type at days 4, 11, 18 and 25 of the graft culture (filled lines). Corresponding hashed lines represent baseline glucose production in PBS controls for each scaffold type. All data represents mean ± s.e.m. from 3 independent jejunal grafts constructed using human colon (green line - hColon), human small intestine (red line - hSI) and piglet small intestine (blue line - Piglet) decellularised scaffolds.

### *In vivo* transplantation of the engineered jejunal grafts

To examine the survival and maturation of the engineered jejunal grafts *in vivo*, we further transplanted the grafts in immunodeficient mice under the kidney capsule or in subcutaneous pockets that present different *in vivo* microenvironments regarding stromal infiltration and vascularisation. First, the jejunal grafts seeded on piglet scaffolds were collected at Day 18 of the culture and transplanted under kidney capsule. Engrafted engineered intestine was collected 7 days after transplantation, which showed signs of neo-vascularisation macroscopically surrounding the scaffold (Fig. 6a). Serial histological sectioning along the length of the scaffold demonstrated the presence of continuous rings of intestinal epithelium that were positive for human nucleoli staining (Fig. 6b,c and Supplementary Fig. 5a). 3D volume reconstruction of the serial histological section data revealed segments of continuous tubular structures maintained throughout the graft, indicating formation of intestinal lumens (Fig. 6d, movie 1). Surprisingly, immunostaining of AB-PAS and ALPI was largely negative in the graft, suggesting a lack of goblet cell and enterocyte differentiation in the engineered graft after engraftment to kidney capsule (Supplementary Fig. 5b). Interestingly, a large population of cells co-expressed with vimentin and αSMA was noted surrounding the lumen, indicating that there is a strong infiltration of host myofibroblasts into the scaffold (Fig. 6e). Immunofluorescent staining further demonstrated high expression of stem cell markers OLFM4 and SOX9 (Fig. 6f,g). Together, the results suggest that transplantation of the jejunal graft in kidney capsule drives the intestinal epithelium to undifferentiated state, which could be due to the strong stromal infiltration that recapitulates the microenvironment of the intestinal stem cell niche.

**Figure 6.**
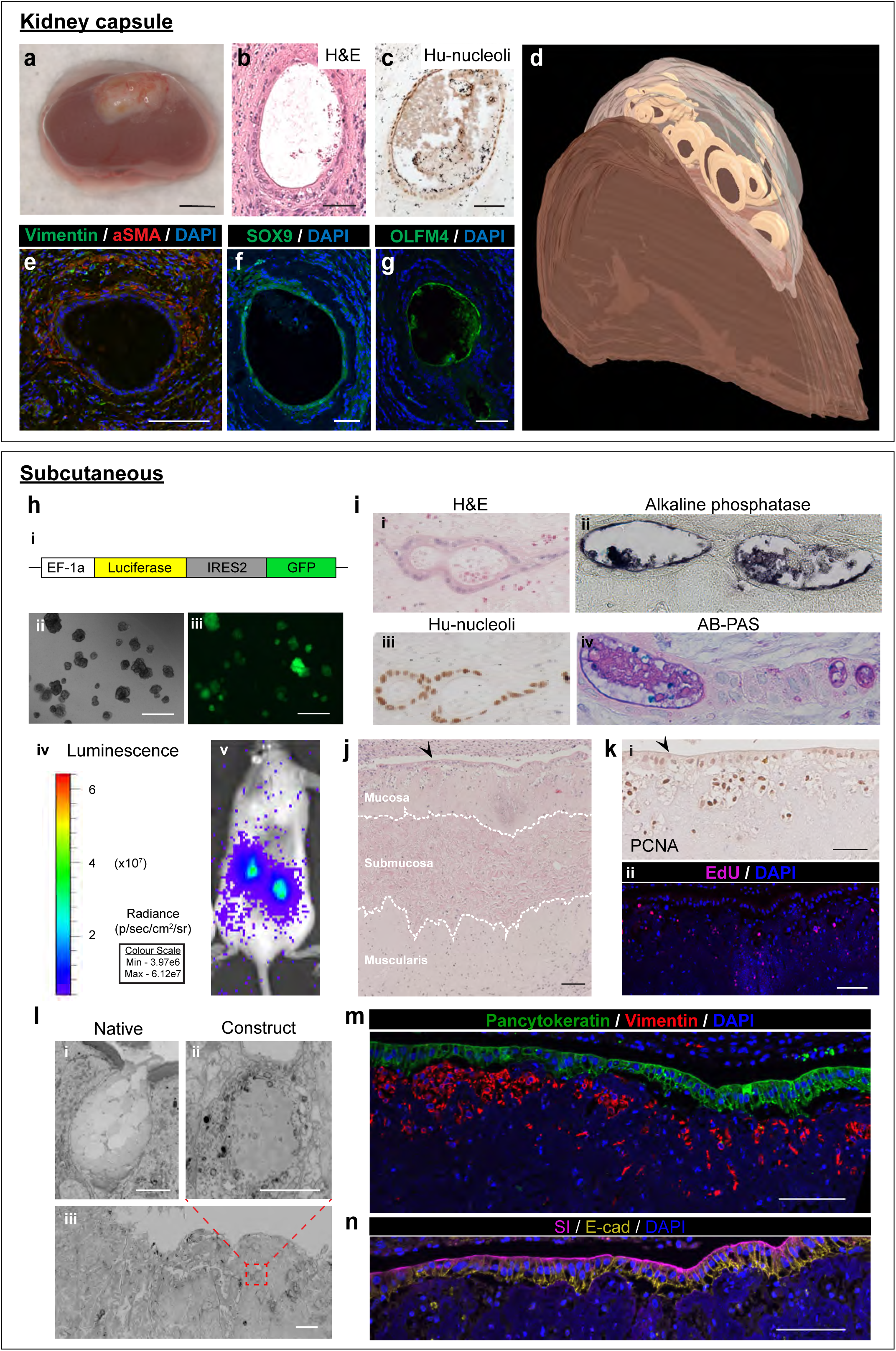
Characterisation of the engineered jejunal graft following *in vivo* transplantation. Histological analysis of the Jejunal grafts transplanted under kidney capsule (a-g) or subcutaneously (h-n) for 7 days. (a) Macroscopic image of the kidney harvested after subcapsular implantation of the jejunal grafts (derived from piglet scaffolds); scale bar represents 2mm. (b,c) Histology of the transplanted jejunal graft was analysed by H&E (b) and human nucleoli staining (c). (d) 3D volume rendered model of the jejunal graft structure transplanted under the kidney capsule. (e-g) Representative immunofluorescent images of the transplanted jejunal grafts using the indicated antibodies. Scale bars represent 50μm. (h) Labelling of the jejunal organoids with luciferase-GFP reporter plasmid for live *in vivo* bioluminescent imaging of the subcutaneous implantation model. Scale bars represent 500μm. (i) Representative histological images of the transplanted jejunal grafts reconstructed using piglet scaffolds using the indicated staining methods or antibodies. Scale bars represent 50μm. (j-n) Histological analysis of the transplanted human jejunal grafts reconstructed using human SI scaffolds using the indicated antibodies. Arrow heads indicate the jejunal epithelium. (l) Electron microscopy analysis identifying mucous granules of goblet cells in the native intestine (i) and jejunal construct (ii,iii). Scale bars represent 20μm. (m,n) Representative immunofluorescent images of the transplanted human jejunal grafts using the indicated antibodies. Scale bars represent 50μm.

Since kidney capsule transplantation was technically challenging for human scaffolds due to the significant thickness of the constructs, we proceeded to test subcutaneous implantation approach that allows engraftment of larger human scaffold constructs. We further pre-labelled the human jejunal organoids with a GFP-luciferase reporter prior to *in vitro* seeding, which enabled us to monitor the epithelial growth *in vivo* with live bioluminescent imaging (Fig. 6h). We first repeated the transplantation experiment using jejunal grafts seeded on piglet scaffolds subcutaneously for comparison. Similar to kidney capsule model, lumens of human nucleoli-positive intestinal epithelial cells were readily detected as soon as 7 days after implantation (Fig. 6i). However, unlike kidney capsule data, ALPI+ enterocytes and AB-PAS+ goblet cells were detected in the graft with significantly less stromal infiltration (Fig. 6i). We postulate that the low stromal infiltration during subcutaneous engraftment may offer a better approach to allow functional differentiation of the jejunal grafts.

Next, we performed subcutaneous transplantation of the jejunal grafts that were seeded on human scaffolds. In order to enhance the growth of ectopically transplanted jejunal grafts in this model, we further investigated the potential use of the drug Teduglutide for tissue engineering applications. Teduglutide is clinically licenced for use in patients with intestinal failure to promote intestinal adaptation by increasing villus height and crypt depth^41^. Since the receptor (GLP2R) of Teduglutide was expressed in both human fibroblasts and organoids (Supplementary Fig. 5c), we decided to treat the mice with Teduglutide during *in vivo* transplantation. Similar to piglet scaffold data, jejunal graft with human scaffold was also able to form lumen after transplantation (Supplementary Fig. 5d). Human SI scaffold was significantly thicker than piglet ones, where the three histological layers (mucosa, submucosa and muscularis) of the graft were clearly preserved after transplantation (Fig. 6j). A distinctive monolayer of human intestinal epithelial cells was formed on the mucosal luminal surface of the implanted scaffold. Proliferating cells (indicated by PCNA and Edu staining) were detected in both epithelial and pericryptal stromal cells of the scaffold (Fig. 6k). Electron microscopy analysis identified mucous granules within the graft epithelial cells, suggesting the presence of early goblet cell differentiation (Fig. 6l). Immunofluorescent staining of human-specific pancytokeratin and e-cadherin confirmed the epithelial identity of the cells (Fig. 6m,n). Expression of human-specific stromal marker vimentin further confirmed the survival of human intestinal fibroblasts in the injected grafts after transplantation (Fig. 6m). Importantly, the jejunal-specific brush border enzyme sucrase isomaltase was broadly expressed in majority of the jejunal graft (Fig. 6n), indicating that human ECM scaffold supports and maintains intestinal epithelial cell differentiation towards enterocyte lineage *in vivo*.

## DISCUSSION

Recent advances of organoid culture have opened up exciting avenues for regenerative medicine^42-45^. However, the regenerative potential of organoids is limited to small lesions only, while larger-scale tissue damage such as organ failure would require more sophisticated approaches. Tissue engineering offers promising alternative treatment strategy for patients with organ failure. However, engineering of complex organs such as intestine is still far from clinical translation due to the limited supportive evidence from pre-clinical studies. Establishment of a robust and relevant protocol for timely organ reconstruction using patient-derived human materials is a critical step prior translation. In the current study, we described a robust protocol to isolate and expand primary hISCs from children with IF. The ability to rapidly derive and expand autologous hISCs and stromal cells from little biopsies is crucial for IF children with very limited residual intestine. We also confirmed that the primary SI organoids retained their region-specific identities and differentiation capacity after pro-longed culture under expansion conditions. Importantly, these primary SI organoids can robustly repopulate the decellularized intestine in customised bioreactors and develop appropriate absorptive and digestive function, which can further survive and mature *in vivo*. To our knowledge, this is the first pre-clinical study that describes protocol for robust generation of primary human SI organoids and decellularized scaffolds from IF patients for jejunal graft reconstruction. Previous work on tissue engineering of small intestine focused mainly on either synthetic scaffolds or ECM scaffolds derived from rodent and porcine^33,37,46-49^. A recent study reported tissue engineering of small intestine using ESC-derived hIOs^38^. The authors showed that the survival and intestinal fate were only maintained when these hIOs were seeded on artificial polyglycolic/poly L lactic acid scaffolds but not on decellularized ECM scaffolds. Their results raised concerns on the clinical application of decellularized ECM scaffold for intestinal tissue engineering. One plausible explanation of the data is that ESC-derived hIOs may not be the ideal cell source for tissue engineering using decellularized ECM scaffold, possibly due to epigenetic plasticity induced during reprogramming. Unlike many other previous tissue engineering studies that focused on iPSCs, synthetic scaffolds or rodent ECM matrix, the use of primary human materials (both cells and decellularized ECM matrix) in this study is a highly relevant step towards clinic translation. Our findings further suggest that routine banking of intestinal epithelial organoids and stromal cells may conceivably be introduced as a new clinical standard at the point of intestinal resections in infants and children undergoing bowel resection (e.g. necrotising enterocolitis) to facilitate personalised intestinal reconstruction in the future.

The DET decellularization protocol represents a more physiological approach for the fabrication of non-immunogenic biological scaffolds as compared to synthetic polymers. The ability to preserve the biological and structural features of the decellularised scaffolds after cryopreservation further suggests the potential of these scaffolds to become “off-shelf” clinical products^13^. Previous studies have demonstrated the conservation of the biological ECM molecules (e.g. collagen and GAGs) in rodent or piglet intestinal scaffolds using targeted methods such as immunohistochemical staining of the tissues^34,37,46^. Here, we characterised the ultrastructure and biochemical composition of the human derived intestinal scaffolds comprehensively and unbiasedly using electron microscopy, Raman spectroscopy and mass spectrometry^50^. Raman imaging demonstrated that the global biomolecular cues amongst the histological layers were largely conserved between the native and decellularized scaffolds. Moreover, spectral signatures arising from the human SI and colon scaffolds were remarkably analogous. This was further supported by the unbiased mass spectrometry analysis that showed highly conserved global and ECM protein profiles despite significant microarchitectural differences at the mucosal layer (i.e. crypt-villus units in the SI scaffold as compared to the lack of villi in the colonic scaffold). Importantly, we demonstrated that human jejunal organoids were able to engraft, proliferate, differentiate and function appropriately on both human SI and colon ECM scaffolds with jejunal-specific brush border enzyme expression and ECM remodelling, suggesting that the intestinal epithelia have cell-intrinsic role in defining the region-specific identity. Our findings support the potential use of colon scaffolds for SI graft reconstruction, which has strong clinical implications for two main reasons. Firstly, colon derived from cadaveric donors or resected in children affected by conditions such as Hirschsprung’s disease could be decellularized, stored, and donated for therapy. Secondly, in conditions such as midgut volvulus, in which typically the large bowel is preserved, there is the potential to convert the IF patient’s own colon to SI by replacing the mucosal layer with jejunal organoids as an alternative treatment solution. Finally, our data generated on piglet scaffolds suggest that porcine ECM matrix could also be used as for human organ reconstruction.

In addition, we further demonstrated that the engineered jejunal grafts were structurally and functionally competent for cell engraftment and vascularisation when transplanted *in vivo*. While the strong stromal infiltration during kidney capsule engraftment resulted in stem cell maintenance and lack of differentiation, grafts were well-differentiated in the subcutaneous implantation model. Intestinal stromal cells, including fibroblasts and myofibroblasts, are known to constitute the essential microenvironment for ECM remodelling and growth factors secretion for stem cell maintenance^51^. The results highlight the essential niche role of stromal cells for the survival and maintenance of the engineered grafts. These outcome can also be enhanced *in vivo* by the drug teduglutide^52^, which was used clinically in IF patients to increase villus height and crypt depth of the intestinal mucosa^53^. Our data showed that administration of teduglutide enhanced the survival of the subcutaneously transplanted intestinal grafts, a site with less host stromal infiltration of the scaffolds. Given that teduglutide is clinically approved for IF patients, we believe that its application during transplantation of the engineered intestinal grafts in IF patients may further promote survival and differentiation of the grafts *in vivo*.

In summary, the current study describes a robust protocol for the timely jejunal mucosal reconstruction using IF patient-derived primary materials including organoid, fibroblasts and ECM scaffolds. The next step will be to scale up the jejunal graft for pre-clinical testing in larger animal models using orthotopic transplantation. Our findings represent an important conceptual advance towards the reconstruction of full thickness of intestinal wall including the enteric neuromuscular cells that drive peristalsis along the length of the engineered intestine for autologous transplantation to IF patients.

## ACKNOWLEDGEMENTS

We thank STEMCELL™ Technologies for providing organoid culture reagent and Pei Shi Chia for coordinating patient information. We also thank the Francis Crick Institute’s Science Technology Platforms: Experimental Histopathology, Electron Microscopy, Mass Spectrometry Proteomics, Mechanical Engineering, Biological Resources Facility and In vivo imaging. This work was funded by the Horizon 2020 grant INTENS 668294 on the project ‘INtestinal Tissue ENgineering Solution for children with short bowel syndrome’. VSWL lab is funded by the Francis Crick Institute, which receives its core funding from Cancer Research UK (FC001105), the UK Medical Research Council (FC001105), and the Wellcome Trust (FC001105). PDC is supported by NIHR Professorship and the Oak Foundation. This research was supported by the NIHR GOSH BRC.

## AUTHOR CONTRIBUTIONS

PDC and VSWL conceived the project. LM, PDC and VSWL designed the study and wrote the manuscript. LM performed the experiments and analysed the data. IM, AK and AB performed in vitro culture and histology analysis. AW, EH, JK and LC performed the experiments on electron microscope and data analysis. RG and GT performed the Raman spectroscopy experiment and data analysis. LM, PF and BS performed the mass spectrometry experiment and proteomic analysis. LM, MO and SE (Eaton) performed the *ex vivo* functional analysis of the engineered intestinal grafts. AFP, AMT and SE (Eli) performed piglet scaffold decellularization experiments. PB performed *in vivo* transplantation experiments. NT coordinated human tissue collection. LM, SE (Eaton), PB, PDC and VSWL critically discussed the data and manuscript.

## METHODS

### Animals and human tissue

Immune deficient NOD-SCID IL-2Rγnull (NSG) female mice, aged 9 - 14 weeks old, were used in all experiments (obtained from the Francis Crick Institute Biomedical Research Facility). All mice were housed in the animal facility at the Francis Crick Institute. All experiments were performed with ethical approval under Home Office Project License PPLs 70/8904 and 70/8560.

Porcine (Sus scrofa domesticus) SI from the ‘Pietrain’ breed was used to derive piglet SI scaffolds. Piglets up to 3 kg in weight were euthanized following criteria outlined by the JSR veterinary advisors. Once sacrificed, the animals were transported to the lab via courier and the intestine was harvested immediately on arrival (within 6 hours of euthanasia).

Ethical approval for the use of human intestinal tissue was obtained from the Bloomsbury NRES committee (REC reference 04-Q0508-79). The Committee was constituted in accordance with the Governance Arrangements for Research Ethics Committees and complied fully with the Standard Operating Procedures for Research Ethics Committees in the UK. Informed consent for the collection and use of human tissue was obtained from all patients, parents or legal guardians at Great Ormond Street Hospital, London. Endoscopic biopsies for organoid isolation were sought from patients with an established diagnosis of intestinal failure on parenteral nutrition. Surgical resections for scaffold fabrication were sought from patients undergoing stoma formation or closure procedures.

### Organoid and fibroblast culture conditions

Endoscopic biopsy specimens were cut into finer pieces and washed in cold PBS. For organoid isolation, the tissue fragments were incubated in 2mmol/L EDTA cold chelation buffer, consisting of distilled water with 5.6mmol/L Na2HPO4, 8mmol/L KH2PO4, 96.2mmol/L NaCl, 1.6mmol/L KCl, 43.4 mmol/L sucrose, 54.9 mmol/L D-sorbitol, 0.5mmol/L DL-dithiothreitol) for 30 minutes at 4°C as previously reported^29^. Following this incubation period, crypts were released from the fragments by shaking vigorously. The supernatant was centrifuged at 800RPM for 5 minutes at 4°C, to form a pellet of intestinal crypts, which were washed in Advanced Dulbecco’s modified Eagle medium (DMEM) / F12 supplemented with 5% penicillin/streptomycin, 10mmol/L HEPES and 2mM of GlutaMAX. The crypts were then resuspended in Basement Membrane Extract^®^ (BME) and seeded in a single droplet on pre-heated 48-well plates. The BME was polymerised for 20 minutes at 37°C, before adding 250μL/well of either human IntestiCult™ Organoid Growth Medium (STEMCELL Technology, #06010) or human organoid basal culture media consisting of conditioned media produced using stably transfected L cells (Wnt 50%; R-spondin 10%; Noggin 5%) and the following growth factors: B12 1X (Invitrogen), Nicotinamide 10mM (Sigma-Aldrich), N-acetyl cysteine 1mM (Sigma-Aldrich), TGF-β type I receptor inhibitor A83-01 500nM (Tocris), P38 inhibitor SB202190 10μM(Sigma-Aldrich), Gastrin I 10nM (Sigma-Aldrich), EGF 50ng/ml (Invitrogen). Rho-kinase inhibitor Y-27632, 10µM was added to the culture media for the first week in culture, at a concentration of 10μM. The media of each well was changed every 2 days. Organoids in expansion were cultured in 3μM CHIR99021. Organoids in differentiation phase were cultured in 10µM DAPT for 48 hours. For the isolation of human intestinal fibroblasts, intestinal fragments left over from the chelation step above were washed in PBS and placed on the bottom of tissue culture dishes with DMEM supplemented with 10% heat inactivated Fetal Bovine Serum (FBS), antibiotics and 1% non-essential amino acids (all from Sigma-Aldrich). Fibroblasts grew from the fragments within 3-4 days. Cells used for seeding experiments were between passages 3-10.

### mRNA isolation and quantitative PCR

RNA was extracted according to the manufacturer’s instructions (Qiagen RNAeasy). cDNA was prepared using High-Capacity cDNA Reverse Transcription Kit (Applied Biosystems, #4368813). Quantitative PCR detection was performed using PowerUp™ SYBR^®^ Green Master Mix (Applied Biosystems, A25742). Assays for each sample were run in triplicate and were normalised to housekeeping genes β-actin, where data was expressed as mean ± s.e.m. Primer sequences are listed in Supplementary Table 4.

### Decellularization of SI and colon

The detergent-enzymatic treatment (DET) for decellularization, previously established on rat small bowel, was adapted for porcine and human intestine^34^. One cycle of DET consisted of the following steps: (i) 24 hours of deionised water at 4°C; (ii) 4 hours of 4% sodium deoxycholate (Sigma) at room temperature; (iii) 1 hour of deionised water at room temperature; (iv) 3 hours of 2000kU DNase-1 (Sigma) in 1M NaCl (for human tissue) or 0.15M NaCl and 10mM CaCl (for piglet tissue) at room temperature^34^. After harvesting piglet small intestine, the mesenteric artery, mesenteric vein and lumen were cannulated using a 29-gauge surgical cannula. The lumen of the intestine and the mesenteric artery were perfused with continuous fluid delivery of DET solutions using a Masterflex L/S variable speed roller pump at 3ml/min, for two cycles. Whole sections of human intestinal tissue collected from surgery were washed in cold PBS containing 5% penicillin/streptomycin to remove luminal contents, before starting decellularization by immersion in DET solutions and gentle agitation on a roller. For human scaffolds, two to three cycles of DET were performed, depending on the weight of tissue received from the patient’s resection. Gamma irradiation at a dose of 16000Gy for 15 hours was applied to sterilise the scaffolds and then preserved at 4°C, in PBS containing 1% penicillin/streptomycin prior to use in cell culture.

### DNA and ECM Quantification

Tissue samples were taken at random immediately post-harvesting and after decellularization protocol for DNA and ECM components quantification. DNA was quantified using a PureLink Genomic DNA Mini Kit (Thermo Fisher). The final concentration of DNA in the samples was measured using a NanoDrop (model NanoDrop 1000 Spectrophotometer by Thermo Fisher). ECM components were quantified using a QuickZyme Collagen assay kit (QuickZyme Biosciences) to measure the collagen, a Blyscan Sulfated 18 Glycosaminoglycan Assay kit (Biocolor) for the glycoaminoglycans (GAGs) and a Fastin Elastin Assay kit (Biocolor) for elastin, according to manufacturers’ instructions.

### Histology

Samples were fixed in 10% Neutralised Buffered Formalin (Sigma) at room temperature overnight, dehydrated in graded alcohols, paraffin embedded and sectioned at 4μm. Tissue slides were stained according to manufacturers’ instructions with Haematoxylin and Eosin (H&E) (Thermo Fisher), Alkaline Phosphatase (Vector Laboratories), Alcian-Blue Periodic Acid Schiff (Sigma). Picrosirius Red (PR), Elastic Van Gieson (EVG) and Alcian Blue (AB) (Thermo Fisher) staining was used to assess retention of collagen, elastin and glycosaminoglycans respectively.

### Immunostaining

For immunofluorescence studies, paraffin embedded slides were rehydrated, permeabilized with 0.3% Triton X100 (Sigma, UK) for 30 minutes at room temperature. Heat mediated antigen retrieval was performed using a sodium citrate buffer (pH6). For whole mount immunostaining of intestinal organoids, cells were fixed with 4% PFA at room temperature for 20 minutes. Primary antibodies were diluted in 1% BSA/PBS/0.01% Triton X-100 and applied overnight at 4°C. Samples were incubated with Alexa Fluor secondary antibodies (Invitrogen) for 45 min at room temperature, washed and mounted with ProLong™ Diamond Antifade Mountant with DAPI (ThermoFisher). EdU staining with the Click-iT EdU Alexa Fluor 568 Imaging kit (Life Technologies) followed the manufacturer’s protocol. DNA was stained with DAPI (Molecular Probes). Images were acquired using a Leica SP5 confocal microscope.

For immunohistochemistry, antigen retrieval was performed using a sodium citrate buffer. Slides were permeabilized using a 0.2% Triton X100 (Sigma, UK) for 30 minutes at room temperature, and blocked with 5% bovine serum albumin (BSA) for 30 minutes. Primary antibodies were detected using peroxidase conjugated secondary antibodies using standard protocols as described previously. Image analysis and capture was performed using a Leica stereomicroscope or an inverted Nikon microscope. All antibodies used are listed in Supplementary Table 5. Images were processed using ImageJ and Adobe Photoshop. Quantifications were performed on raw images however for presentational clarity, adjustments of brightness and contrast were applied equally to all panels of a given figure.

### Mass spectrometry and data analysis

Four biological replicates of both decellularized colon and SI ECM scaffolds were used for mass spectrometry analysis. Samples were prepared for analysis as previously reported^54^. Briefly, 1mg of lyophilised decellularised tissue was solubilised in 8M urea containing 10mM dithiothreitol. Iodoacetamide was added to a final concentration of 55mM and incubated for 30 minutes at room temperature protected from light. Proteins were treated with PNGaseF (25,000 units/mg) overnight. An initial protein digest using Lys-C (10 µg/mg for 4 hours) was followed up with two successive tryptic digests (20 µg/mg overnight; 10 µg/mg for 4 hours). All enzyme reactions were performed at 37⍰C. Four biological replicates each of colon and intestine scaffolds were processed. Peptide material was cleaned up using C18 Sep-Pak columns (Waters 50 mg sorbent, WAT054955). Eluates were dried in a speedvac concentrator. Dried peptides were solubilised in 0.1 % trifluoroacetic acid (TFA) to a concentration of approx. 5 µg/µl, then diluted to 0.25 µg/µl in a glass auto-sampler vial immediately prior to analysis. Each of the eight samples were analysed in technical triplicate (approx. 1 µg per injection) using a ThermoFisher Scientific QExactive mass spectrometer coupled to an UltiMate 3000 HPLC system for on-line liquid chromatographic separation. Each sample was initially loaded onto a C18 trap column (ThermoFisher Scientific Acclaim PepMap 100; 5 mm length, 300 µm inner diameter) then transferred onto a C18 reversed phase column (ThermoFisher Scientific Acclaim PepMap 100; 50 cm length, 75 µm inner diameter). Peptides were eluted at a flow rate of 250nL/min with a stepped gradient of 5-25% buffer B (80% acetonitrile, 0.1% formic acid, 5% DMSO) for 60 minutes followed by 25-40% for 20 minutes. Higher energy Collisional Dissociation (HCD) was used for MS/MS peptide fragmentation. Singly-charged and unknown charge state precursor ions were not analysed. Full MS spectra were acquired in the orbitrap (m/z 300–1800; resolution 70k; AGC target value 1E6) with the MS/MS spectra of the ten most abundant precursors from the preceding MS survey scan then acquired (resolution 17.5k, AGC target value 1E5; normalised collision energy 28 eV; minimum AGC target 1E2). Selected precursors were dynamically-excluded for 15 s.

Raw data files were processed using MaxQuant software (version 1.6.0.13) for protein identification and quantification using intensity based absolute quantification (iBAQ). iBAQ values were calculated for colon and SI by combining technical and biological replicates. A SwissProt Homo sapiens protein database (downloaded July 2017 containing 26,389 reviewed sequences) was searched with a fixed carbamidomethylation of cysteine modification and variable oxidation of methionine and protein acetylation (N-term) modifications. Protein and peptide false discovery rates were set at 1 %. The MaxQuant protein groups output file was imported into Perseus software (version 1.4.0.2) for further statistical analysis and data visualisation. Common contaminant proteins and reverse sequences were removed. Intensity values were log2 transformed and Gene Ontology cellular compartment (GOCC) descriptions were added by Perseus. Protein detection was called when it was detected in at least 3 out of the 4 biological replicates of either colon or SI scaffolds. This resulted in 377 total proteins detected in these intestinal scaffolds (Supplementary Table 2).

### Raman Spectroscopy

Raman imaging was conducted using a Renishaw RA816 Biological Analyser coupled to a 785 nm laser excitation source that is reshaped using cylindrical lenses to produce a line illumination (Renishaw plc, Wotton-under-edge, UK). A total laser intensity of approximately 158mW was focused onto the sample through a 50×/NA 0.8 objective. A 1500 l/mm grating was used to disperse the laser light providing a spectral range of 0 to 2100 cm-1 in the low wavenumber range. The RA816 series undergoes a fully automated calibration and optimization sequence to ensure optimal performance, including calibration to the 520.5 cm-1 silicon peak. Raman imaging was conducted on colonic and small intestine sections previously embedded in paraffin wax and cut at 8 µm. Sections were mounted onto stainless steel slides, deparaffinised in xylene and rehydrated in graded alcohol and distilled water prior to Raman analysis. A total of thirty single point spectra were acquired from each histological region of both the large and small intestine (mucosa, submucosa and muscularis propria) using a 15 seconds integration time. Large Raman maps were acquired using the Renishaw StreamLine™ mode using a 4.4 µm step size and 1.1 seconds integration time per pixel. A total of 8,544 spectra were acquired across the colon map and 22,825 spectra across the small intestine map.

Prior to any analysis all spectra were pre-processed to remove all non-chemical effects of the data acquisition process. All spectra were truncated between 400 – 1800 cm^-1^ which encompasses the fingerprint region where the majority of all biological signal lies. Cosmic ray removal was conducted using the width of feature and nearest neighbour methods in Renishaw’s WiRE 5 software. Spectra were then imported into MATLAB R2017a (MathWorks, Natick, MA, USA) where baseline correction to a third order polynomial was conducted using the modified polyfit method^55^. Spectra were vector normalised and then analysed in both WiRE 5 and MATLAB R2017a.

Point spectra obtained from each distinct histological site in the colon and small intestine (mucosa, submucosa and muscularis propria) before and after decellularisation were averaged and plotted on the same axis. The most obvious differences pre- and post-decellularisation were highlighted. Principal component analysis (PCA) was then carried out to ascertain whether the biochemical difference between each distinct intestinal layer post-decellularisation could be observed. This is an unsupervised multivariate analysis technique that allows an effective reduction in the dimensionality of the spectral dataset and hence facilitates the identification of combinations of highly correlated variables that best describe the variance in the data. Both of these procedures were carried out in MATLAB R2017a. The large high spatial resolution Raman maps were analysed in WiRE 5 (Renishaw plc, Wotton-under-Edge, UK) using unsupervised Multivariate Curve Resolution – Alternating Least Squares (MCR-ALS) approach for initial exploratory analysis to determine the global composition of each specimen. Using the reconstructed component curves, we were able to identify some of the components abundant within each histological layer. Direct Classical Least Squares Component analysis was then used to acquire component images of a number of known biomolecules using previously acquired reference spectra. This enabled us to ascertain the spatial distribution of known biomolecules within the full thickness of the intestinal wall.

### In vitro scaffold seeding using bioreactor system

Sterilised sections of acellular intestinal scaffolds were immobilised in custom made mini platforms and placed at the bottom of 24 well tissue culture plates. When seeding human scaffolds, human jejunal fibroblasts were trypsinised and resuspended in DMEM before seeding by multiple microinjection via 26g cannulae (Terumo SKU:SR+DU2619PX) into the lateral walls of the scaffolds, at a density of 1×10^5^ cells/cm^2^. Three days after fibroblast injection, human jejunal organoids were trypsinised and seeded onto the mucosal surface of the scaffolds at a density of 1×10^6^ cells/cm^2^. The cells were allowed to engraft for a period of 30 minutes at 37°C, before covering the whole scaffold with basal human intestinal organoid culture media. The scaffolds were maintained for another 4 days in static culture conditions, before transferring the scaffolds into perfusion plates (Amsbio #AMS.AVP-KIT-5) and connecting this to a bioreactor circuit. The bioreactor circuit consisted of a media reservoir (custom made by Chem Glassware UK Manufacturers Ltd, London) with a 0.22μ air filter, inlet and outlet tubing (Cole Parmer cat.no. 224-2081), a peristaltic pump (Cole Parmer cat.no. 224-1505) and 3 way stopcocks (Becton Dickinson UK Ltd cat.no. 394601) for media sampling at both inlet and outlet points of the circuit. Cultures were maintained for 14 days in dynamic culture conditions, with media circulating at a rate of 3ml/min. Piglet scaffold seedings were seeding in a similar manner, without pre-injections of fibroblasts due to the lack of accuracy of injection in the thin scaffold walls.

### β-Ala-Lys-AMCA peptide update assay

At day 4, 11, 18 and 25, the grafts in culture (or unseeded scaffolds as controls) were transferred from the bioreactor circuit into a 6 well plate and rinsed several times with PBS. They were then incubated with fresh human organoid culture media containing 25µM of β-Ala-Lys-AMCA for 2 hours at 37⍰C. After incubation the media was removed and the graft was rinsed in cold PBS three times. The grafts (or unseeded scaffolds as controls) were fixed in 4% paraformaldehyde for 30 minutes at room temperature. Whole mount immunostaining was performed as described above, staining with Alexa Fluor 568 phalloidin to mark cellular boundaries. The samples were then imaged using a Leica SP5 inverted confocal microscope. The fluorescence signal of β-Ala-Lys-AMCA was acquired using the UV laser.

### Citrulline detection assay

Graft culture supernatants were collected at baseline, day 11 and 25. Citrulline levels were quantified by spectrophotometry according to methods reported previously^56^. Briefly, 20µl of each test sample was added to 20µl of water and 10µl of working urease solution, then incubated at 37⍰C for 30 minutes. 150µl of chromogenic reagent was then added to the solution and incubated for a further 60minutes at 100⍰C to allow colour development. Absorbance was read at 520nm in a 96 well microtitre plate using a Tecan microplate reader (Infinite^®^ M1000 PRO). Concentration was determined by comparison to a citrulline standard curve.

### Disaccharidase enzyme activity assay

Grafts in dynamic culture were transferred from the bioreactor circuit into 6 well plates. The scaffolds were washed in PBS three times then incubated with a solution of 56mM sucrose in PBS (or PBS alone in control wells) for 60 minutes. Aliquots of the solution were then sampled for glucose detection using the Amplex™ Red glucose/glucose oxidase assay kit (ThermoFisher cat.no. A22189) according to manufacturer’s protocol. Briefly, 50µl of the reaction working solution was added to 50µl of the test samples in a 96 well black flat bottom microtitre plate in triplicates, and incubated in the dark for 30 minutes at room temperature. The fluorescence (excitation 535nm, Emission 590nm) was measured using a Tecan microplate reader (Infinite^®^ M1000 PRO). Concentration was determined by comparison to a glucose standard curve.

### Electron microscopy and micro-CT imaging

For routine SEM imaging, decellularised human scaffold samples were fixed in 2% glutaraldehyde in 0.1 M phosphate buffer and kept at 4°C for 24 hours. Samples were then washed with 0.1 M phosphate buffer and cut into segments of approximately 1 cm in length and cryoprotected in 25% sucrose, 10% glycerol in 0.05 M PBS (pH 7.4) for 2 hours, then fast frozen in nitrogen slush and freeze fractured at −160°C. Samples were then returned to the cryoprotectant solution and allowed to thaw at room temperature. After washing in 0.1 M phosphate buffer (pH 7.4), the material was fixed in 1% OsO4 in 0.1 M phosphate buffer (pH 7.4). After rinsing with distilled water, specimens were dehydrated in a graded ethanol-water series to 100% ethanol, critical point dried from CO2 and mounted onto aluminium stubs using sticky carbon tabs. Samples were coated with a thin layer of Au/Pd using an ion beam coater (Gatan UK). Images were recorded using a Jeol 7401 field emission gun scanning electron microscope.

For routine TEM, MicroCT imaging, serial block face SEM imaging and montage SEM imaging, recellularised scaffold grafts were fixed overnight at room temperature in 10% neutralised buffered formalin (Sigma) followed by a second fixation step in 2.5% glutaraldehyde/4% paraformaldehyde in 0.1 M phosphate buffer (pH7.4). The sample was post-fixed in 2% reduced osmium (4% OSO4, 1.5% K3FE(CN)6) for 60 minutes on ice and then washed in ddH20.

To check the orientation of cells on recellularised scaffolds, the sample was embedded in CYGELTM (BioStatus, Leicestershire, UK) in an Eppendorf tube and a microCT scan was performed at 4kV/3W, with no filter, 1601 projections and a pixel size of 7.3379 µm using an Xradia 510 Versa (Zeiss). The data was automatically reconstructed using Scout-and-Scan™ Control System Reconstructor software (Zeiss) and viewed in TXM3DViewer software (Zeiss). With the presence of cells confirmed, the sample was immersed in ice, the CYGEL™ washed off the sample with ddH20 and the sample trimmed to approximately 1 mm^3^ blocks. The trimmed blocks were then incubated in 1% aqueous thiocarbohydrazide at room temperature for 20 minutes then washed in ddH20. The blocks were incubated in 2% aqueous osmium tetroxide for 30 minutes at room temperature and washed in ddH20. This was followed by a further incubation in 1% aqueous uranyl acetate at 4⍰C overnight. The blocks were washed in ddH20 and then incubated in Walton’s lead aspartate for 30 minutes at 60⍰C before being dehydrated through a graded series of ethanol, infiltrated with Durcupan resin (Sigma-Aldrich) and polymerised for 48 hours at 60⍰C.

For routine TEM, 70 nm sections were cut on a UC6 ultramicrotome (Leica), picked up on formvar-coated slot grids and imaged in a Tecnai G2 Spirit Biotwin (ThermoFisher) with an Orius CCD camera (Gatan UK). For Serial Block Face SEM, Samples were trimmed to the region of interest, mounted on an aluminium pin (Leica Microsystems) and sputter coated with 2 nm of platinum. See Supplementary Table 6 for serial block face SEM imaging conditions. For SEM montage images, after the orientation of the cells within the block was determined by microCT, 200 nm sections were cut on a Powertome ultramicrotome (RMC) and picked up on ITO-coated coverslips. The coverslips were mounted on SEM pin stubs (Agar Scientific) using a sticky carbon tab and sputter coated with 0.5 nm platinum. Sections were viewed in a Quanta FEG scanning electron microscope (ThermoFisher) using a backscattered electron (BSE) and large montage images acquired using MAPS 1.1 software (ThermoFisher). See Supplementary Table 7 for SEM montage imaging conditions. The montaged image in Figure 5 C i was generated by stitching together individual images using TrakEM2, a plug-in of the FIJI framework^57^. The montaged image in Figure 5 C ii and 5 C iii were generated by stitching together individual images using MAPS 1.1 software (ThermoFisher).

### Lentivirus preparation for human organoid labelling

The lentiviral vector pHIV-LUC-ZsGreen (Addgene Inc. MA, USA, Plasmid #39196, kind gift from Dr Bryan Welm, Department of Surgery, University of Utah) was used to generate a lentivirus containing both ZsGreen fluorescent protein and firefly luciferase from an EF1-alpha promoter. Human jejunal organoids were labelled by lentiviral transduction as previously reported39. Briefly, LUC-ZsGreen lentivirus was produced by co-transfecting 293T cells with the above plasmid along with packaging vectors PAX2 and VSV-G envelope plasmid (kind gifts from Dr Hans Clevers). Transfection was performed according to manufacturer’s instructions for 8⍰hours at 37°C. The medium (Opti-MEM^®^) was exchanged for virus collection. After 24 hours, the virus-containing medium was purified by centrifugation at 2500⍰rpm (4⍰°C) and filtered through a 0.45μm membrane and ¼ volume of PEG-itTM was added to the filtered supernatant before ultracentrifugation at 2300 x⍰g for 30⍰mins at 4°C (SW28 rotor, Optima LE80K Ultracentrifuge, Beckham). The viral pellet was resuspended in 1ml of pre-cooled Opti-MEM^®^ (Gibco), aliquoted and stored at −80⍰°C. Human organoids were dissociated into single cells and cultured in the presence of 250µl 2x organoid culture media and 250µl viral particles and 2.5µl of TransDuxTM. Transduction efficacy was determined measured as the proportion of cells expressing the fluorescent protein ZsGreen 72⍰h after transduction.

### Bioluminescence imaging (BLI)

BLI was performed using an IVIS Spectrum in vivo imaging system (PerkinElmer, Waltham, MA, USA) and Living Image 4.3.1 software (PerkinElmer). Mice were injected intraperitoneal with 150 mg/kg D-luciferin (PerkinElmer) twenty minutes prior to imaging. All images were taken at field of view C or D, with automatic exposure time, pixel binning set to 8, f-stop 1 and open emission filter. This generated pseudo-coloured scaled images overlaid on grey scale photographs, providing 2-dimensional localization of the source of light emission. All images were analysed using Living Image 4.3.1 software (PerkinElmer). Regions of interest (ROI) were drawn manually and the light emission was quantified in photons s-1. ROI shapes were kept constant between images within each experiment.

### *In vivo* transplantation models

NSG mice were anaesthetized with a 2-5% isoflorane: oxygen gas mix for induction and maintenance. The dorsum of each animal was shaved and the skin cleansed with 70% ethanol and povidone-iodine solution. For kidney capsule transplantation (n=3), each seeded graft (cultured as 1cm^2^ patches) was carefully cut in half and gently folded to maintain the epithelial surface internally, before immediately inserting the graft under the capsule of the kidney. For subcutaneous transplantation (n=12), closed blunt scissors were used to create subcutaneous pockets bilaterally in the dorsum of each mice and one folded graft segment was inserted in each pocket. Subcutaneous Teduglutide or vehicle (PBS) was administered at a dose of 0.2 mg/kg/daily after implantation. The mice were sacrificed at 7 days for analysis. 2 hours prior to culling, each mouse was administered a dose of EdU (3ul/g of a stock solution of 10mg/ml).

### 3D volume rendering of kidney capsule transplant data

Serial sections were cut of the paraffin embedded sample. Odd numbered slides were stained with H&E while even numbered slides were kept unstained for further immunostaining analyses. Odd numbered H&E slides were then serially scanned (Olympus VS120 slide scanner) and the region of interest was aligned manually using Amira Software (ThermoFisher). Using the aligned slices, the kidney, scaffold, epithelial ring and lumen were segmented manually and saved as four separate label fields before generating 3D surfaces. A movie was created using Amira Animation Director.

### Statistical Analysis

The n value reported in the manuscript for each analysis/assay represents the number of biological replicates. Data are expressed as mean⍰±⍰SEM. Significance was determined by one-way ANOVA and Tukey’s multiple comparison test; ANOVA with post-hoc Bonferroni test and two-tailed unpaired Student’s t-test. A p-value of less than 0.05 was considered statistically significant. Statistical analysis was performed using GraphPad Prism 6 (GraphPad Software).

## Figure legends

**Supplementary Figure 1.**
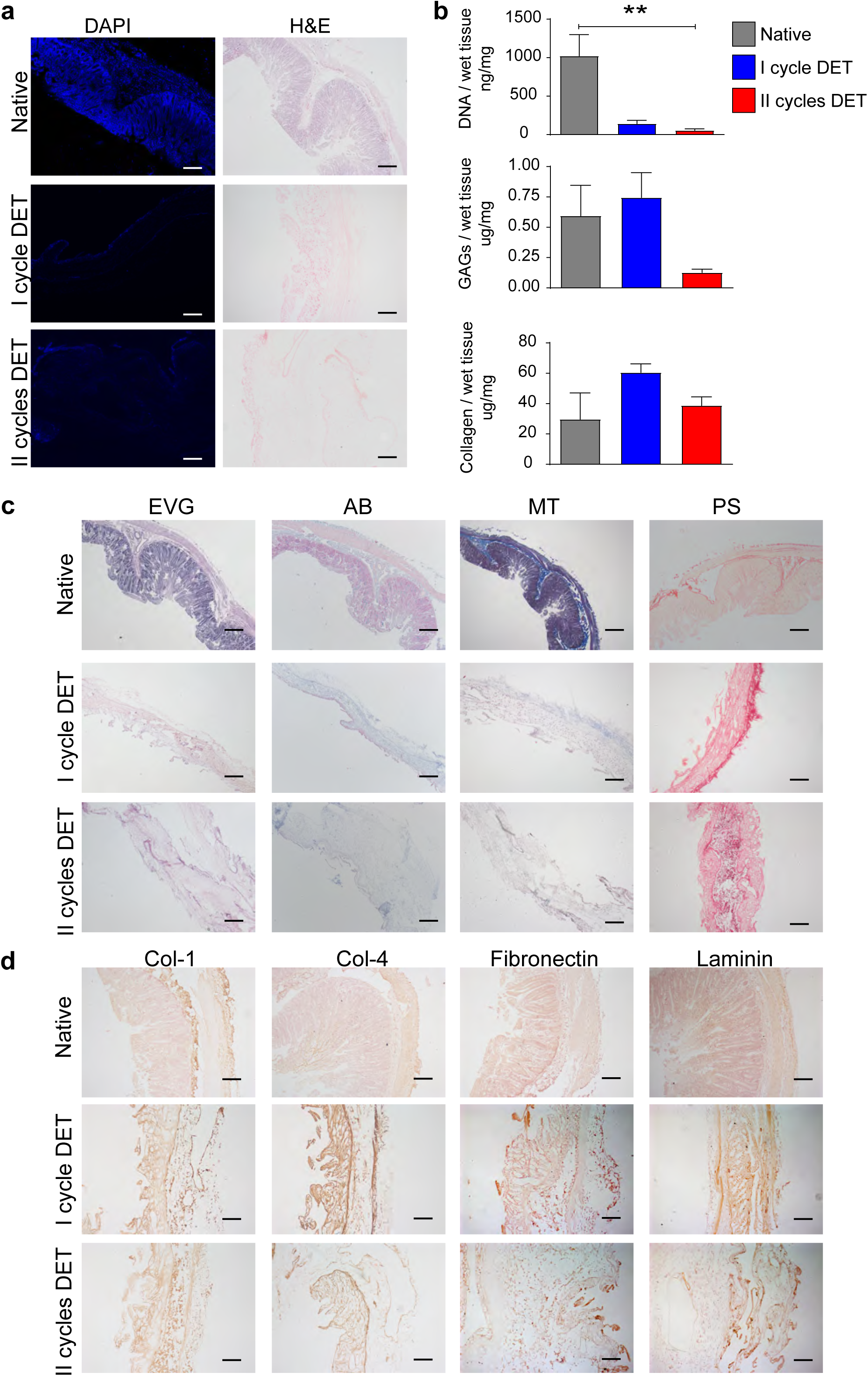
Piglet scaffold characterisation. (a) Representative images of H&E and DAPI stainings of the native piglet intestine and following one and two cycles of DET. Scale bars represent 100µm. (b) Quantification of DNA, glycosaminoglycans (GAGs) and Collagen per milligram of wet tissue in native piglet intestine and following one and two cycles of DET. Data represent mean ± s.e.m using 3 biological replicates. The experiment was repeated 3 times. (c) Representative images of Elastic Van Gieson (EVG) and Alcian blue (AB) stainings confirm preservation of elastin and GAGs respectively following one and two cycles of DET. Representative images of Masson’s trichrome (MS) and Picro-sirius red (PS) stainings confirms maintenance of connective tissue and collagens following one and two cycles of DET. Scale bars represent 200µm. (d) Representative images of immunohistochemical staining for Collagen I, Collagen IV, Fibronectin, Laminin indicating the preservation of these ECM proteins in the scaffold following two cycles of DET. Scale bars represent 100µm.

**Supplementary Figure 2.**
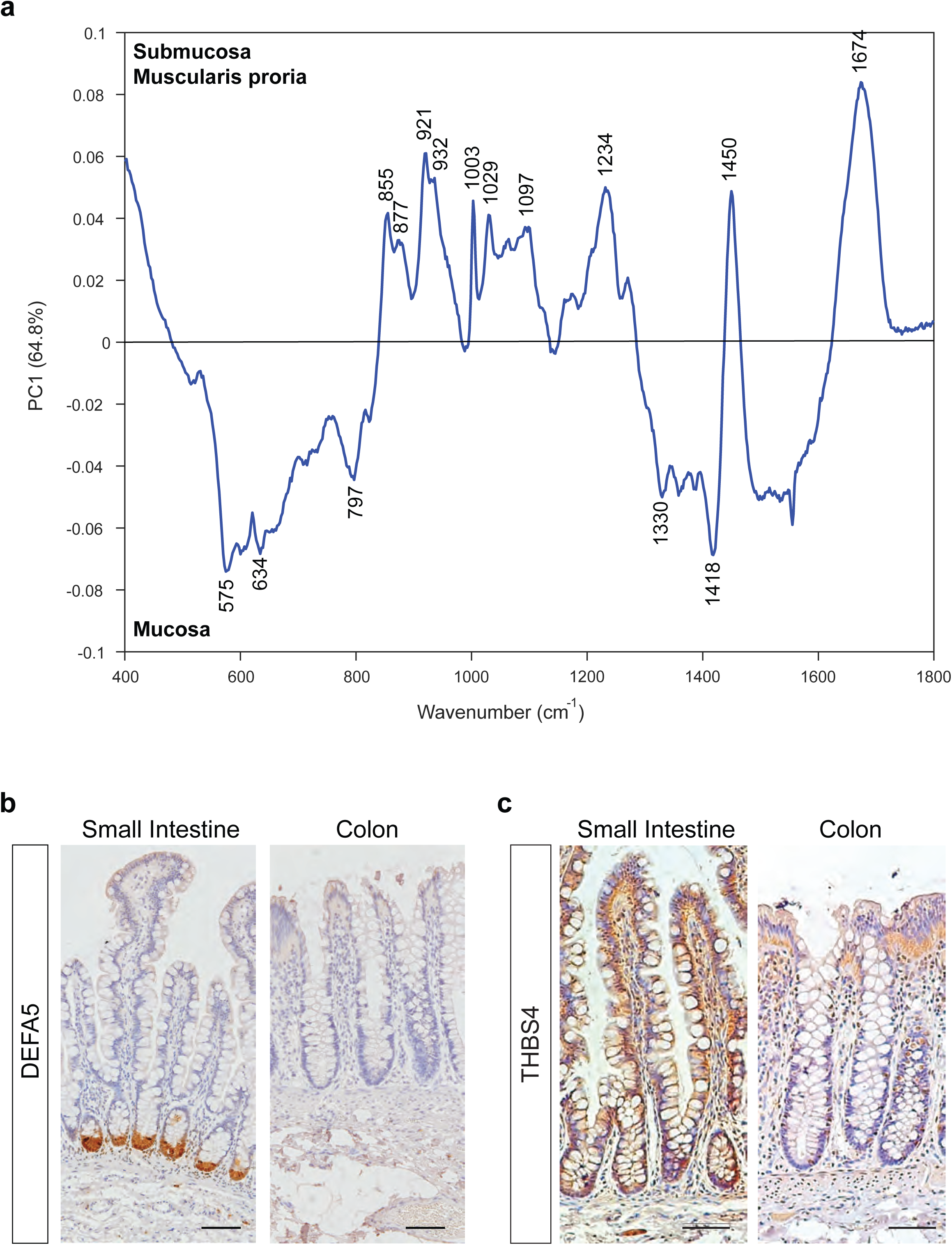
Characterisation of human decellularized intestinal scaffolds. (a) PC1 loading plot associated with the PC1 vs PC2 scores plot shown in Fig. 3c. The spectral features show the distinct biochemical differences that facilitate the differentiation of the mucosal region of the SI and colon from the remaining intestinal layers (submucosal and muscularis propria). The intensity of the corresponding peaks found within the loadings plot indicate their influence on the separation in the scores plotted along the associated axis. (b,c) Representative immunohistochemical staining of the native paediatric SI and colon tissue using the indicated antibodies. Scale bars represent 100μm.

**Supplementary Figure 3.**
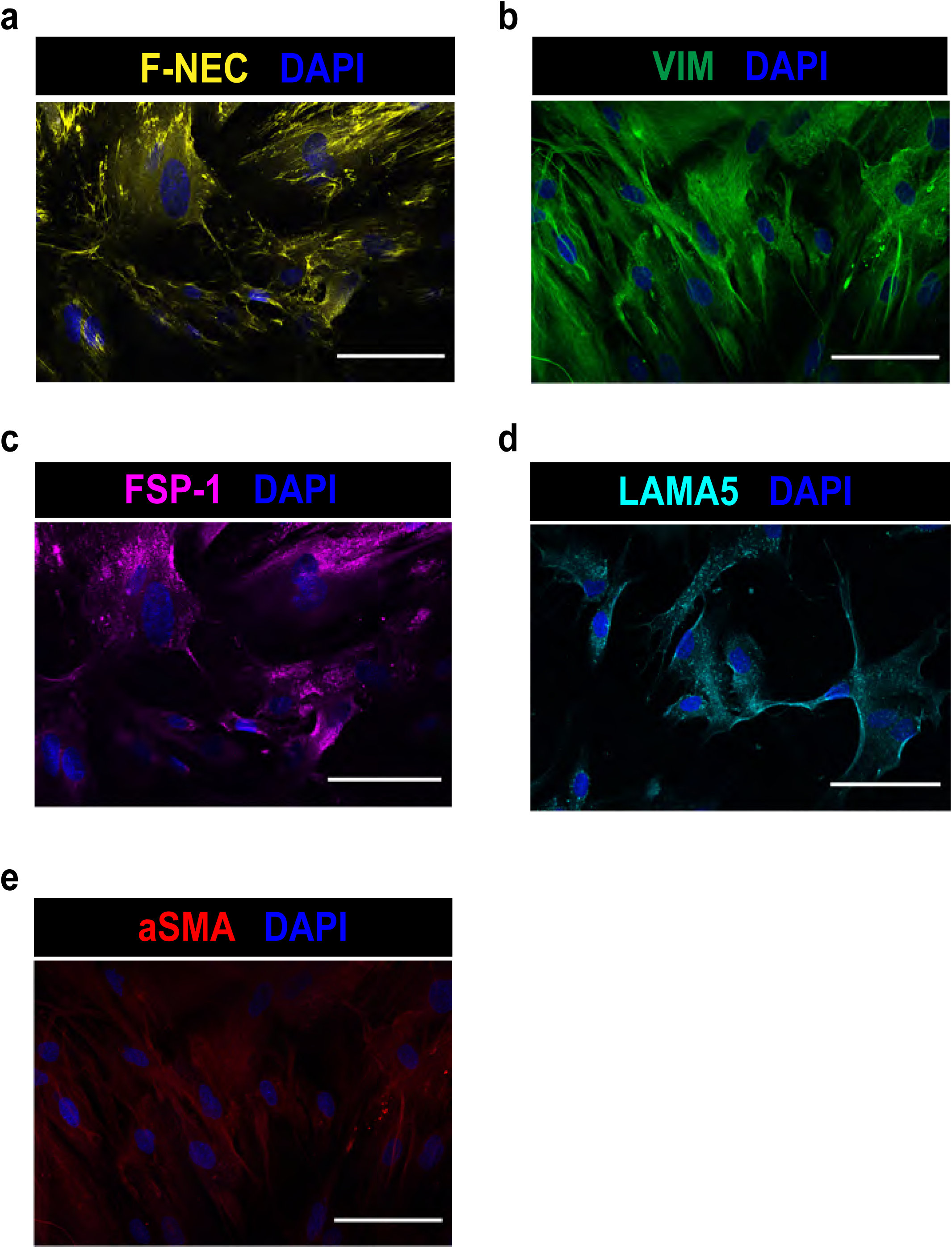
Characterisation of primary fibroblasts derived from human intestinal tissues. Representative immunofluorescent images of primary human SI fibroblasts showing fibronectin (a), vimentin (b), fibroblast surface protein marker-1 (c), laminin alpha 5 (d) and alpha-smooth muscle actin (e). Scale bars represent 50μm.

**Supplementary Figure 4.**
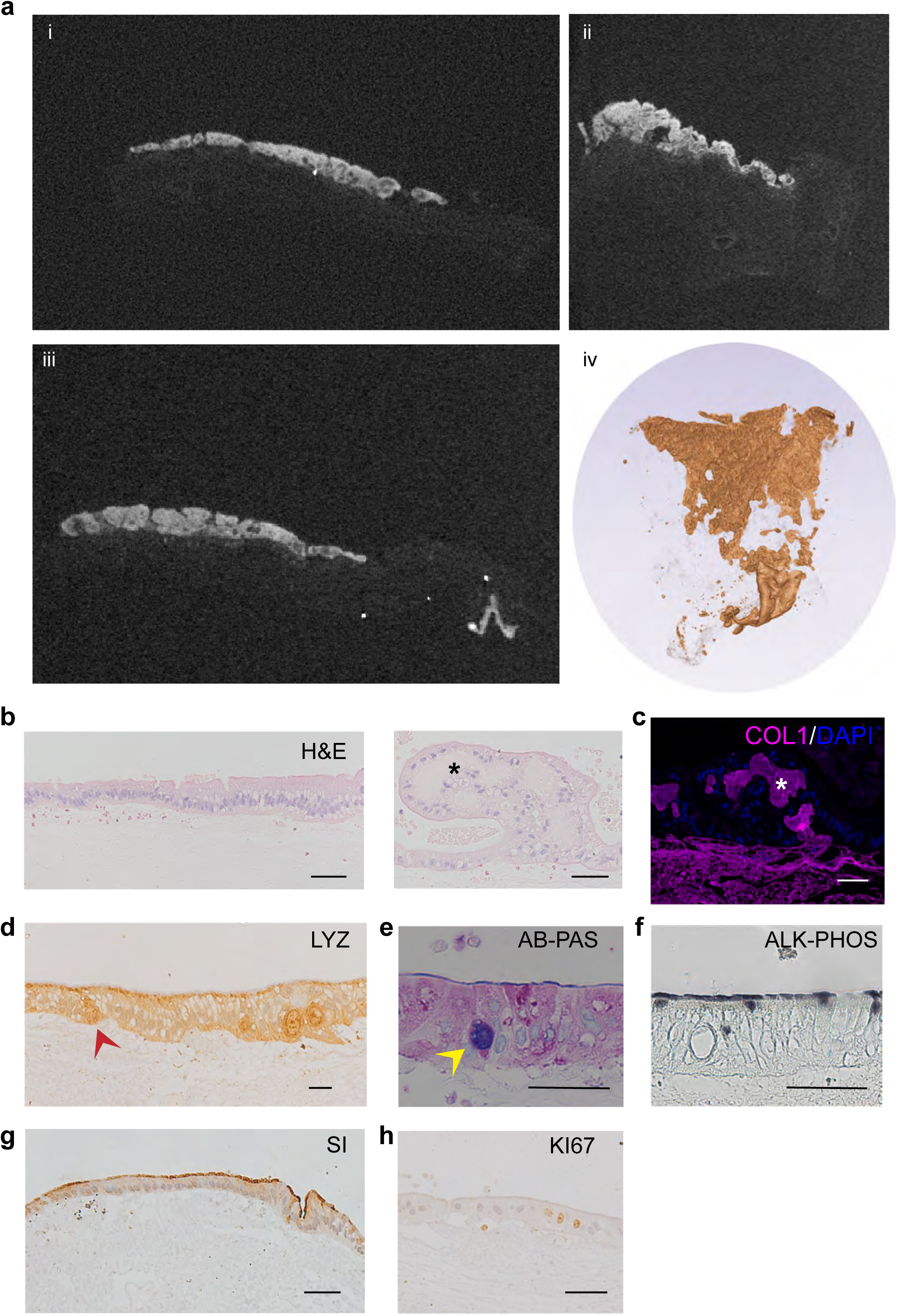
Characterisation of the human jejunal graft reconstructed on piglet scaffold. (a) Micro CT virtual slices of jejunal grafts constructed using piglet scaffolds, showing the jejunal epithelial layer (bright white) on the piglet scaffold (i-iii). 3D volume rendering of the epithelium showing nearly full surface coverage (iv). (b) H&E staining of jejunal graft showing regions of columnar epithelial monolayers as well as regions of new matrix deposition (black asterisks). (c) New matrix deposition is indicated by immunofluorescent staining of collagen-1 (magenta) co-stained with DAPI (blue). (d-h) Representative immunohistochemical images of the engineered jejunal grafts using the indicated antibodies. All scale bars represent 50μm.

**Supplementary Figure 5.**
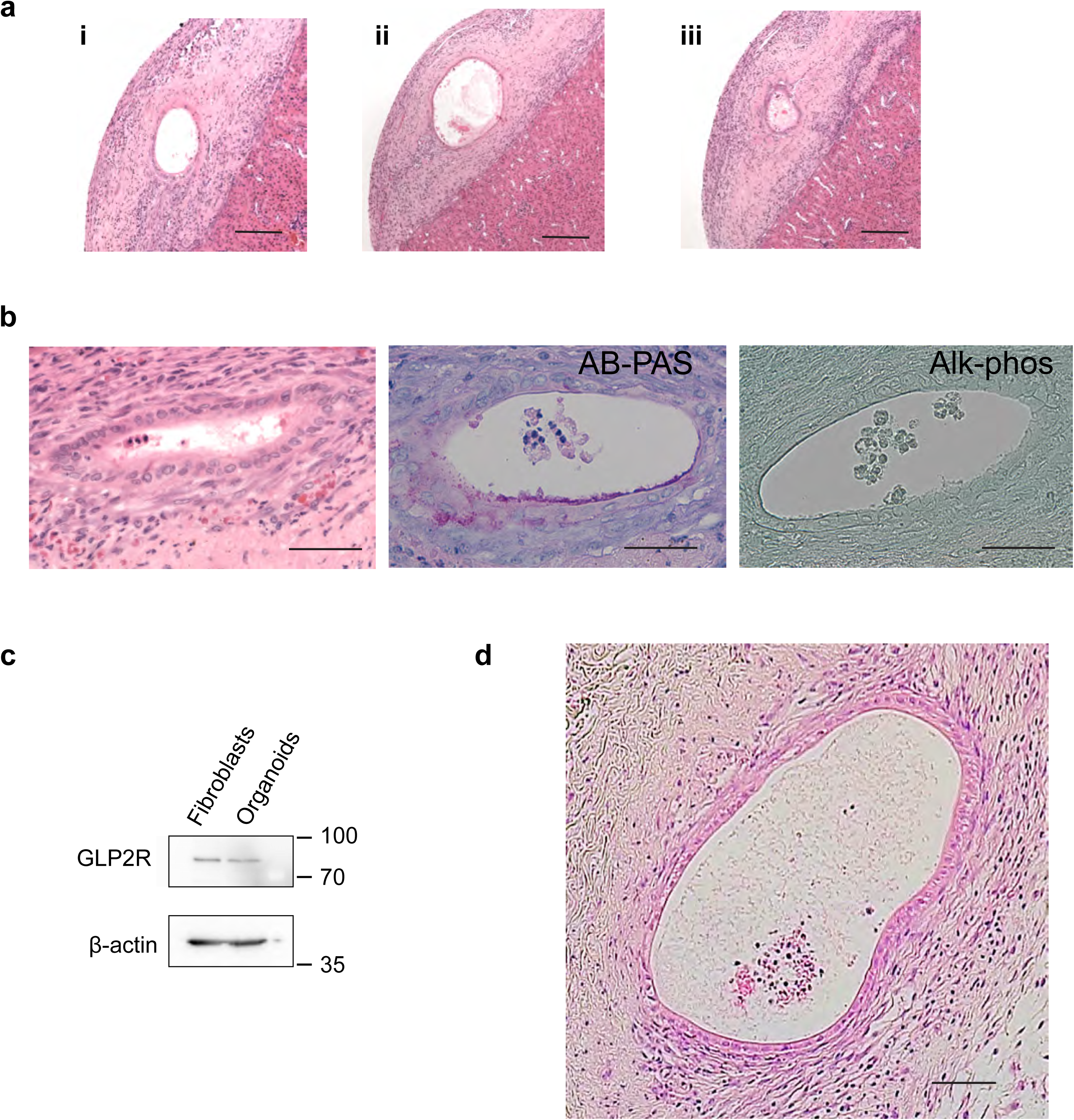
Histological characterisation of the kidney capsule transplanted graft. (a) Serial sections and H&E staining of the jejunal graft following 7 days transplantation *in vivo* under the kidney capsule. (b) Ring of jejunal graft stains negatively for goblet cells (Alcian Blue - Periodic Acid Schiff) and enterocyte brush border marker (Alkaline Phosphatase). All scale bars represent 50μm. (c) Western blot analysis confirming the expression of the GLP2R in human small intestinal fibroblasts and human jejunal organoids when co-cultured *in vitro*. (d) H&E staining of the jejunal graft seeded on human scaffolds *in vivo* under subcutaneous implantation. Scale bars represent 100μm.

